# Evidence accumulation from experience and observation in the cingulate cortex

**DOI:** 10.1101/2025.02.13.638172

**Authors:** Ruidong Chen, Setayesh Radkani, Neelima Valluru, Seng Bum Yoo, Mehrdad Jazayeri

**Author notes:** Equal contribution (Ruidong Chen, Setayesh Radkani).

## Abstract

We use our experiences to form and update beliefs about the hidden states of the world. When possible, we also gather evidence by observing others. However, how the brain integrates experiential and observational evidence is not understood. We studied the dynamics of evidence integration in a two-player game with volatile hidden states. Both humans and monkeys successfully updated their beliefs while playing the game and observing their partner, though less effectively when observing. Electrophysiological recordings in animals revealed that the anterior cingulate cortex (ACC) integrates independent sources of experiential and observational evidence into a coherent neural representation of dynamic belief about the environment’s state. The geometry of population activity revealed the computational architecture of this integration and provided a neural account of the behavioral asymmetry between experiential and observational evidence accumulation. This work lays the groundwork for understanding the neural mechanisms underlying evidence accumulation in social contexts within the primate brain.

## Introduction

A hallmark of cognition is the ability to infer the hidden causes of our experiences. Waking up with an upset stomach, you might wonder if it is due to food poisoning or if you caught the flu. Your first clues come from your symptoms—nausea versus fever would hint at different causes. But you may also rely on others’ experiences. If coworkers recently had the flu, you might lean in that direction, but if your dinner partner has the same symptoms, you may conclude it’s food poisoning. While the capacity to integrate experiential and observational evidence to infer the hidden causes of ambiguous observations is unequivocal, the neural mechanisms that enable such sophisticated computations are not well understood.

The neurobiology of inference and evidence-based decision making in experiential settings has been heavily studied. Among various brain regions, the anterior cingulate cortex (ACC) is thought to play a central role. ACC carries signals related to outcome history, performance monitoring, action and strategy selection, and beliefs about latent action-outcome associations and contexts (*1–21*). Notably, ACC representations persist over relatively long time scales (*22*) and integrate information across events and experiences (*3*, *9*, *10*, *23*). These findings coupled with perturbation experiments and behavioral analysis of subjects with compromised or lesioned ACC (*10*, *15*, *24–26*) have provided strong evidence that ACC encodes behaviorally-relevant beliefs about latent causes in the environment.

We know far less about the computational and neural basis of observational inference and learning but evidence is accumulating. Within the reinforcement learning framework, one approach has been to examine the neural signatures of observed reward and punishment (*27*) in the amygdala, striatum, and many cortical areas (*28–39*). Among these, ACC has been a prominent region of interest that is sensitive to reward given to a conspecific, robustly encodes vicarious outcomes, and is required for vicarious reinforcement and observational fear conditioning (*33*, *35*, *36*, *39–42*). Therefore, ACC may play a more general role in belief updating than spans both experiential and observational settings.

However, most observational studies have relied on relatively simple tasks that either do not require inferring latent causes from observations or do not require integration of experiential and observational evidence. As a result, the neural substrates and mechanisms of social evidence integration remain poorly understood. Furthermore, it is unclear whether there are parallels—or distinctions—between the neural representation of experiential and observational evidence, and how these two sources of information are integrated to form beliefs about the latent state of the world.

Here, we tackle these questions using a combination of behavioral experiments in humans, neurophysiology experiments in nonhuman primates, and neural network modeling. We developed a two-player game in which players had to update their beliefs about an underlying hidden state based on integration of evidence garnered from experience and observation. The behavioral results revealed a familiar asymmetry between experiential and observational evidence (*43–45*) in both humans and monkeys. Neurophysiology recordings in ACC enables us to directly compare the representation of evidence from experience and observation, characterize how they are integrated, and how they impact participants’ beliefs and behavior on a trial-by-trial basis. We found that ACC integrates independent representations of evidence from self-experience and observation into a coherent population pattern of neural activity encoding the dynamic state belief. Moreover, the organization of the population activity associated with self-experience, observation, and integrated belief accounted for the behavioral asymmetry.

## Results

### Behavioral task and performance

We designed a two-player game for humans and monkeys to investigate the behavioral and neural signatures of updating beliefs in the presence of both experiential and observational evidence (Figure 1A-G). Each trial consists of two stages. In the first stage, the players use their respective joysticks to independently choose between a left and a right arena (Figure 1A). They are free to choose either arena (Figure 1D) but this choice is consequential because only one of the arenas may eventually lead to a reward (Figure 1C). Human participants additionally reported their confidence in their choice privately on an increasing scale from 1 to 4.

**Figure 1.**
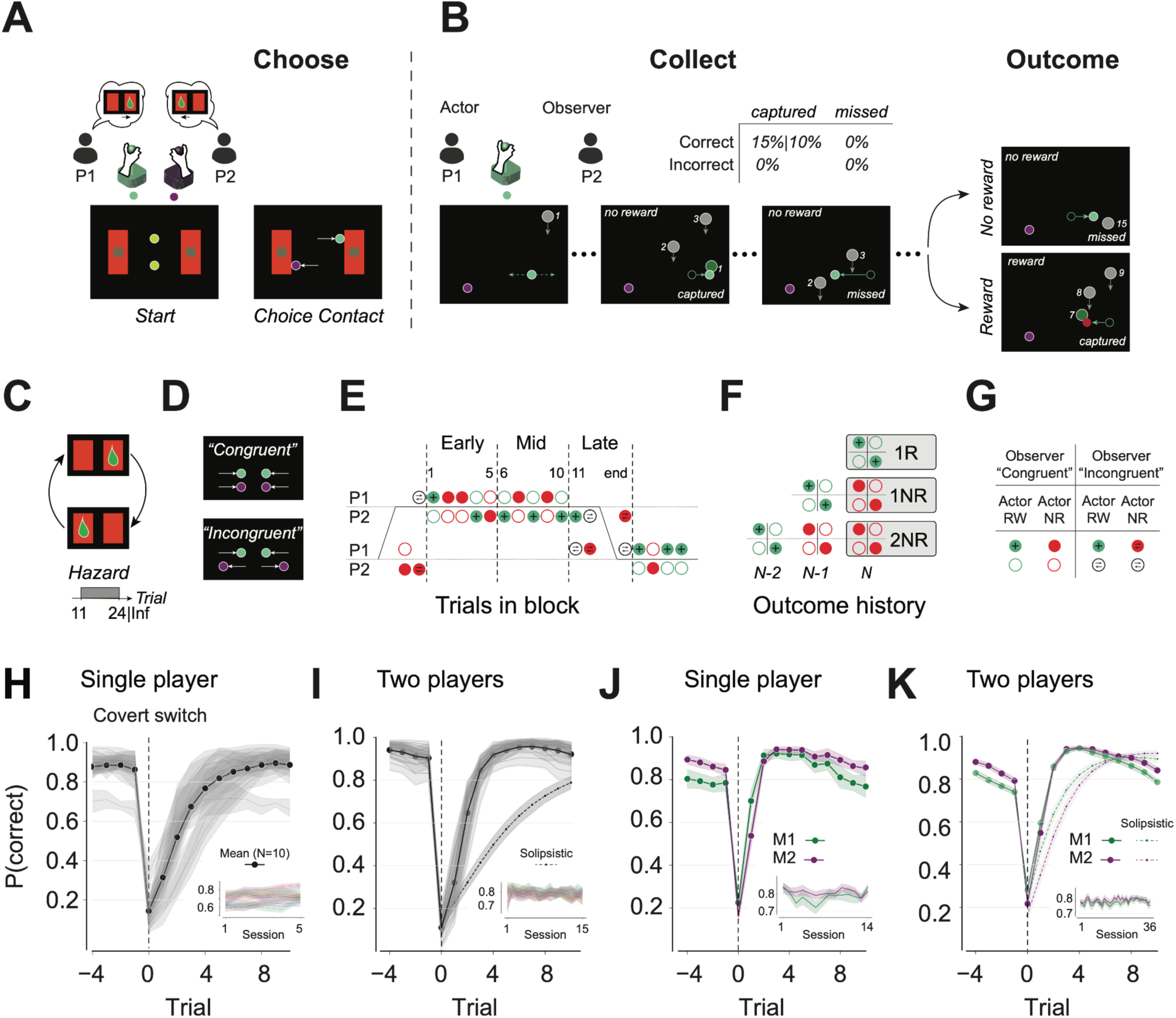
Multi-agent evidence integration task and performance. (A-C) Example trial. (A) Phase 1. In this phase, animals choose one of the two arenas (“Choose”). Start: two avatars (yellow circles) are presented in the middle of two arenas (red rectangles). Choice contact: The avatars’ colors change indicating which avatar belongs to which player (P1: green; P2: magenta) and prompting players to use their joystick to move their avatar toward one of the two arenas based on their belief (think cloud). When an avatar contacts an arena, the choice is registered and the avatar stops moving. The trial proceeds after choices for both animals are registered. Human players additionally indicate their confidence in their choice after making a decision. The players’ choices are revealed, whereas the confidence ratings remain private. (B) Phase 2. In this phase, one of the animals plays a token collection game (“Collect”) for reward (“Outcome”). At the beginning of this phase, P1 and P2 are designated as Actor and Observer, respectively (this designation is randomized across trials). The designation is evident from the placement of the avatars: Observer is placed closer to the bottom of the screen compared to the Actor and below their chosen arena. Immediately after Actor/Observer designations, reward tokens begin to drop from the top of the screen. The Actor must use its joystick to move its avatar sideways and collect as many tokens as possible. The Observer’s avatar remains static on the chosen side throughout the remainder of the trial. The trial terminates either by the delivery of the reward or after a total of 15 tokens have fallen without reward. Outcome: Actor receives reward probabilistically after capturing each token (with probability 0.1 for humans; 0.15 for monkeys) only when they choose the correct arena (table above). (C) Schedule for covert switches. On each trial, only one arena is associated with a non-zero reward probability (green drop). The reward-bearing arena switches covertly (curved arrows) in a blocked fashion. Block switches are governed by a constant hazard after 11 trials. For monkeys, the block length is capped at 25 trials. (D) In the first stage, players can choose the same arena (“Congruent”) or different arenas (“Incongruent”). (E) Example run of trials. The schematic marches through trials from left to right. Two horizontal gray lines correspond to the two arenas (top: left arena; bottom: right arena). Correct arena transitions in a blocked fashion as depicted by a piecewise black line. In this example, the correct arena goes from right to left (bottom to top), stays left for the length of the block (stays on top), and then transitions to right (back to bottom). Disks represent Actor/Observer designations, left/right decisions, and rewarded/unrewarded trials for each player (Actor/Observer: filled/open; left/right: near top/bottom horizontal line; rewarded/unrewarded: green/red; incongruent Observer: opposite arrows). Numbers indicate trial in block starting at the first rewarded trial (i.e., subjective block switch). Trials 1-5, 6-10, and beyond 11, demarcated by dashed lines, are Early, Mid, and Late subdivisions within a block. (F) Nomenclature for outcome history conditions. 1R: congruent rewarded trial. 1NR: congruent unrewarded trial that follows 1R trial in the same arena. 2NR: congruent unrewarded trials that follow 1NR trial in the same arena. The colored disks indicate the combination of Actor/Observer trials that comprise each condition. (G) Key for Actor/Observer designations and rewarded/unrewarded trials for panel E and F. (H) Proportion of trials human participants made the correct choice, P(correct), in the first stage of the single player condition. Trials are grouped relative to covert block switches (trial 0). Shade: 95% confidence interval. Black line shows the average across participants. Inset: P(correct) over sessions, each line corresponds to one participant. Data in the main plot is average across 5 sessions as shown by the range in the inset’s abscissa. (I) Same as H for human participants in the two-player condition. Solid line: Mean performance of all participants. Dashed line: mean predicted performance of the participants had they ignored all observer trials. The performance in the two-player condition was significantly better than predicted by the solipsistic model (p<0.001, t-test). (J) Same as H for the two monkeys in the single player condition. (K) Same as I for the two monkeys in the two-player condition. The performance in the two-player condition was significantly better than predicted by the solipsistic model (p<0.001, t-test).

In the second stage, one player is randomly designated as the actor. The actor controls an avatar in their chosen arena with the joystick and must capture tokens falling from the top of the screen, aiming to collect as many as possible to maximize expected reward. Receiving reward is probabilistic and depends on both the arena and the number of captured tokens during the second stage (Figure 1B). If the actor selects the correct arena, each captured token has a fixed probability of yielding a reward (0.1 for humans and 0.15 for monkeys), and the delivery of reward would terminate the trial (i.e., no more token capture allowed). In the other arena, capturing tokens never results in a reward. Trials without rewards end after all 15 tokens have dropped. The correct arena switches in a blocked fashion. Block switches occur with a probability of ⅓ after a minimum of 11 trials (Figure 1C). From the moment the actor begins to collect tokens to the end of the trial, the other player, whom we refer to as the observer, watches the actor play and witnesses the outcome without receiving any reward.

We designed the experiment to match the sensory experiences of the actor and observer as closely as possible. Specifically, both players view the entire sequence of events across both stages, eliminating informational asymmetries between the actor and observer. Additionally, the designation of the actor happens after both players select their preferred arena, ensuring players report their belief about the correct arena. Finally, because actor/observer designations are random, events on every trial are equally informative to both players for their selection of the arena in subsequent trials. In sum, we ensured that players had all the information needed for making rational inferences about the reward-bearing arena. Matching these external factors is crucial for uncovering asymmetries in evidence accumulation between experienced and observed outcomes due to internal factors.

We collected data from ten human participants (5 fixed pairs) and two monkeys. All participants learned the task in a single-player version before moving onto the two-player version. In the single-player version, choice performance in the first stage was 71.85±7.02% (mean±SD; 5 sessions, Figure 1H) across human participants, 78.39±3.34% in M1, and 80.96±1.65% in M2 (14 sessions, Figure 1J). Next, we quantified performance in the two-player condition. The average choice performance was 79.15±3.05% (mean±SD) across human participants (Figure 1I), 78.68±2.07% in M1, and 80.38±1.73%, in M2 (Figure 1K). Similar to the single-player condition, choices closely followed block switches (Figure 1H-K, S2C-F). In both versions, performance was high before switches, dropped precipitously in the first trial after a covert switch, and recovered within the first few trials of the new block. These patterns are consistent with participants using experiences and observations to update their belief about the correct arena. To further test if participants were attentive when designated as Observer, we used the single-player data to construct a hypothetical agent that ignores trials when designated as Observer (see Methods). The performance of these solipsistic agents (humans: 50.72±10.90%; M1: 74.82±3.27%; M2: 71.40±3.09%) was significantly worse than the performance in the two-player sessions (Figure 1I,K). Consistent with this, observer monkeys’ eye gaze stayed in the active arena during token collection (proportion of time with eye gaze on the active arena as observer, M1: 58.0±22.5%, M2: 87.4±18.7%, Figure S1H,K).

### Humans and monkeys integrate experiential and observational evidence rationally

To evaluate the degree to which participants played the game rationally, we compared their behavior to that of an optimal model (see Methods), which we refer to as the oracle (Figure 2). Specifically, we analyzed the decision to switch on the next trial as a function of three factors: (1) history of trial outcomes (including the current trial), (2) position of the current trial in the block, and (3) number of captured tokens in the current trial. To ensure a fair comparison between actor and observer, we first focus on the subset of trials in which the two players chose the same arena, which we refer to as congruent trials. Later, we discuss the results of the incongruent trials.

**Figure 2.**
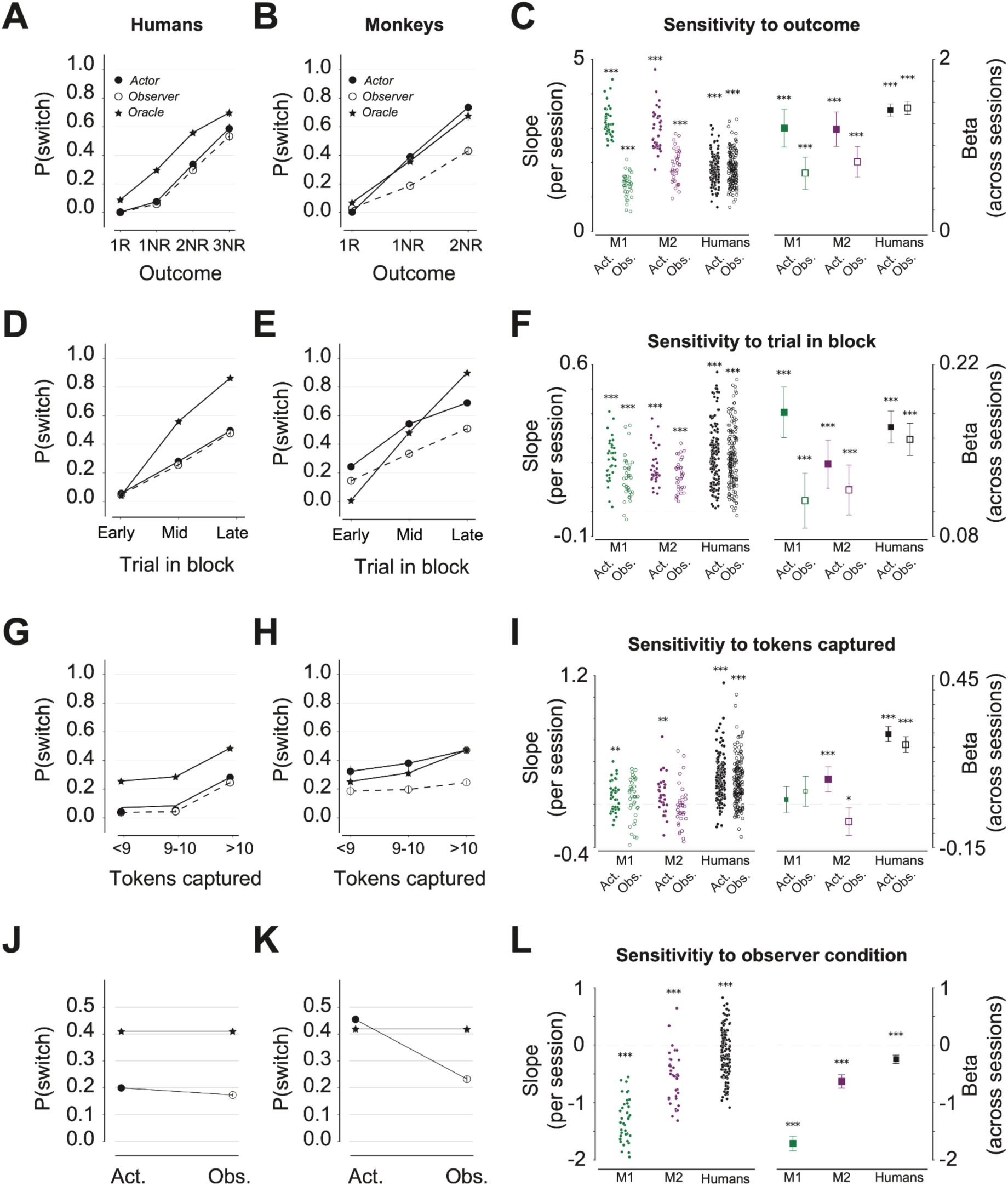
Behavioral characteristics of experiential and observational learning in monkeys and humans. (A-B) Switch probability, P(switch) as a function of trial outcome for congruent trials across humans (A) and monkeys (B). Results are shown separately for Actor (filled circle), Observer (open circle), and Oracle (star). 1R: rewarded trials. *n*NR: *n-th* consecutive unrewarded trial within the same arena. (methods). (C) Left: Regression slope relating P(switch) to outcome history, separately for Actor (filled) and Observer (open) for each participant (two colors) for each session. Stars indicate statistical significance (two-sided t test; ***:P<0.001, **:P<0.01, *:P<0.05). Right: beta values of a multivariable logistic regression relating P(switch) to outcome history, trial in block, and number of captured tokens for Actor (filled) and Observer (open) trials for each participant across sessions. Vertical lines indicate 95% confidence interval; stars indicate statistical significance (two-sided t test; ***:P<0.001, **:P<0.01, *:P<0.05). (D-F) Same as (A-C) for the relationship between P(switch) and trial position in the block for congruent unrewarded trials. (G-I) Same as (A-C) for the relationship between P(switch) and the number of tokens captured for congruent unrewarded trials. (J-L) Same as (A-C) for the relationship between P(switch) and player identity (Actor versus Observer) for congruent unrewarded trials. Regressions include both Actor and Observer trials.

#### Behavioral consequences of outcome history (*Figure 2A-C*)

Since only one of the arenas contains reward, the oracle interprets reward as strong evidence that the arena was chosen correctly. Conversely, unrewarded trials furnish evidence for a block switch, and consecutive unrewarded trials on the same side strengthen this evidence (star, Figure 2A,B). In accordance with the oracle, for both humans and monkeys, the probability of switch, denoted P(switch), was relatively small after rewarded trials and increased monotonically with the number of consecutive unrewarded trials on the same side (Figure 2A-B, S4A-B). This result was robust and evident at the level of single sessions (Figure 2C, left). We further quantified these results using a more conservative logistic regression analysis to account for other factors (see Methods). The regression slope relating P(switch) to the number of unrewarded trials was significantly positive for both humans and monkeys regardless of actor/observer designations (Figure 2C, right). Finally, an analogous multiple-regression analysis of confidence reports in human participants corroborated this finding (Figure S3A)

#### Behavioral consequences of trial position in the block (*Figure 2D-F*)

Since covert switches are less likely early on compared to later in the block, the oracle is more likely to switch later in the block (star, Figure 2D,E). One consideration for testing this prediction is that neither the oracle nor the participants were aware of block switches. To address this point, we inferred subjective block switches from the behavior (Figure S2E-F) and registered trial position within subjective blocks (see Methods). We grouped trials to three approximately equal bins, denoted Early, Mid, and Late, and quantified how switch behavior changed across the bins. In accordance with the oracle, P(switch) for both humans and monkeys increased monotonically with trial position in the block (Figure 2D-E, S4C-D). This effect was evident in single sessions and survived a more conservative logistic regression analysis regardless of actor/observer designations (Figure 2F). Confidence reports in human participants corroborated this finding (Figure S3A).

#### Behavioral consequences of the number of captured tokens (*Figure 2G-I*)

In the second stage, the number of captured tokens was lawfully related to difficulty, suggesting that participants understood that they must collect as many tokens as possible (see Methods; Figure S2G-H). Since each captured token in the correct arena is associated with a non-zero reward probability, the oracle treats unrewarded trials with larger numbers of captured tokens as stronger evidence for a block switch (star, Figure 2G-H). Qualitatively, this effect was evident in human participants’ behavior (Figure 2G, S3A), and to a lesser degree in monkeys (filled, Figure 2H, S4E-F). However, a more rigorous multiple regression analysis indicated that the effect was significant only for human participants, and in Actor condition for one of the animals (Figure 2I, S3B).

#### Behavioral consequences of the choice incongruence (Figure S5)

We also analyzed participants’ behavior in the incongruent trials (trials in which participants chose opposite arenas). Overall, incongruent trials were more frequent later in the block (Figure S5A,D), which is consistent with higher confidence in choice early in the block. Participants reported lower confidence following incongruent trials (Figure S5C). Moreover, across human participants and for one of the monkeys, the actor was more likely to switch following incongruent trials (Figure S5B,E-G), which is consistent with players being influenced by one another’s decisions on top of their experiences and outcomes.

### Humans and monkeys discount observational evidence

One notable feature of our experimental design was the symmetry of experience between the actor and observer: the actor designations were random and sensory experiences across the two players (except from the direct receipt of juice reward in monkeys) were identical. Because of this symmetry, the oracle evaluates trial outcomes identically across actor and observer designations (star, Figure 2J-K). In contrast, humans and monkeys weighted observational evidence less than experienced evidence (Figure 2J-K). This asymmetry was significant for both humans and monkeys but stronger in monkeys, possibly because monkeys – not humans – received juice reward. This asymmetry was evident at the level of single sessions (Figure 2L, left), and survived a more stringent regression analysis of actor and observer trials with the observer condition as an additional factor (Figure 2L, right). This result indicates that humans and monkeys were less responsive to unrewarded trials as an observer. Notably, this asymmetry was not due to players spending less time looking at the screen when designated as Observer (Figure S1E-M). This asymmetry was also evident in humans’ confidence reports when they chose to stay in the same arena as the preceding trial (Figure S3A).

### Experiential and observational evidence integration in the cingulate cortex

Our behavioral results provided compelling evidence that monkeys, like humans, integrate experiential and observational evidence to infer latent state switches. To investigate the underlying neural substrates and computations, we recorded neural activity in ACC (see Table S1 for stereotaxic coordinates) simultaneously from the two animals as they played the game.

Our recordings (M1: 31 sessions; M2: 20 sessions; simultaneous recording in 19 sessions) yielded 1628 units (M1: 859; M2: 769). Many ACC neurons were task-modulated and carried signals related to various observable and latent task variables throughout the trial (Figure 3). During the first decision stage, some neurons were unmodulated (Figure 3A, bottom left), some were sensitive to the choice of arena, and some were differentially modulated for congruent and incongruent trials (Figure 3A, bottom right). After actor/observer designations when the actor began collecting tokens, modulated neurons were sensitive to conjunctions of player identity (actor versus observer) and the choice of arena (Figure 3B, bottom). Finally, many neurons were sensitive to trial outcome with a mixture of selectivity depending on player identity (Figure 3C, bottom).

**Figure 3.**
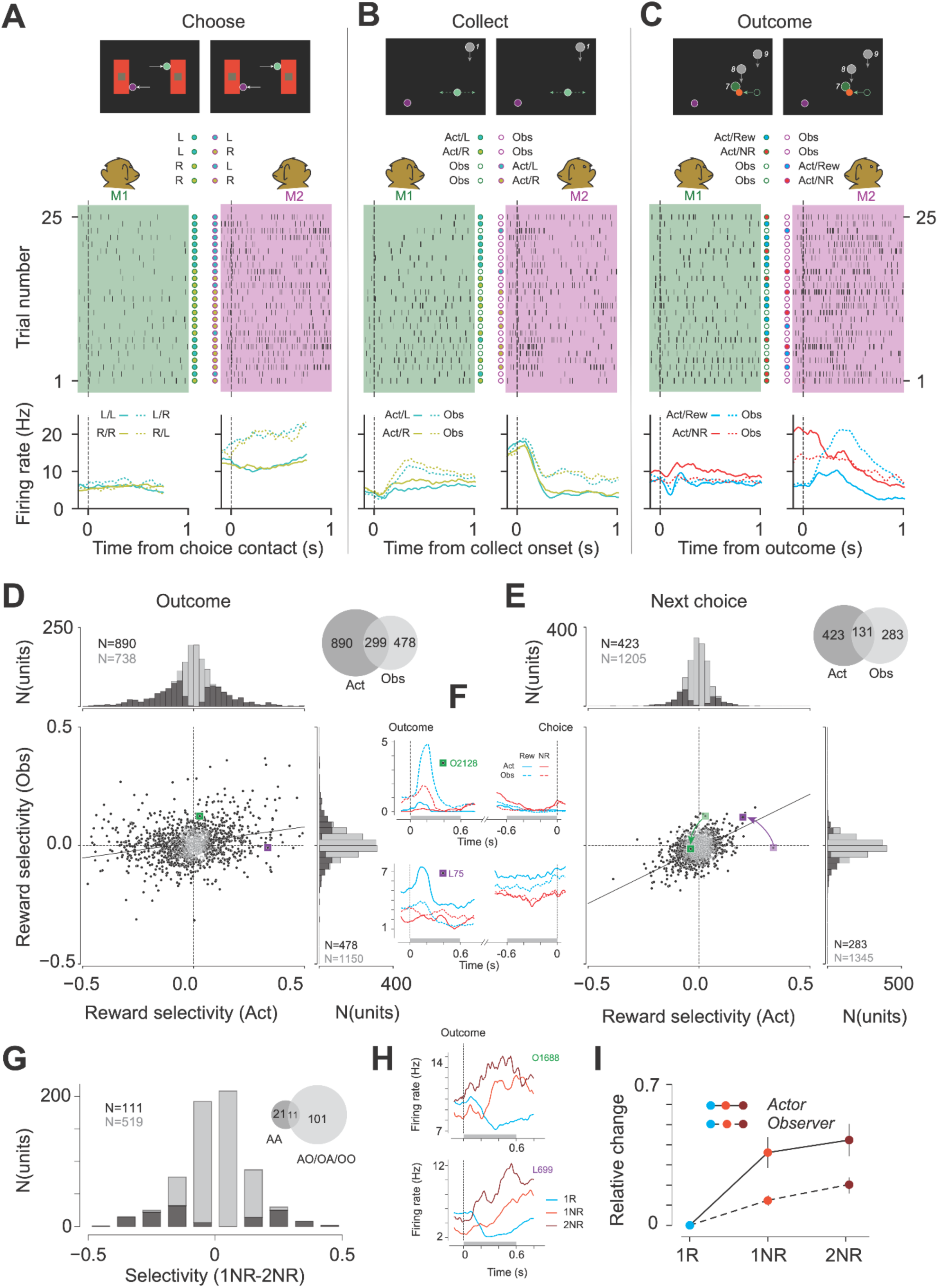
ACC neurons encode and integrate actor and observer outcome. (A) Choice phase. Top: screen display for two monkeys at the moment when they have both made contact with the choice rectangles. L and R refer to the choice of each monkey. Filled circles indicate actor, open circles indicate observer. Middle: raster plot of action potentials recorded from a unit in M1 (left, green shade) and M2 (right, magenta shade). Dashed vertical line: the time of choice contact of the monkey’s own avatar. Trials are chronologically ordered from 1 to 25. Bottom: firing rate histogram of each unit conditioned on animal choices. The unit in M2 has a higher firing rate on trials when choices differ. (B) Same as (A) with the same two units for the collect phase. Dashed vertical lines indicate the onset of token collection. (C) Same as (A) with the same two units for the outcome phase. Dashed vertical lines indicate the cue for reward or end of trial. (D) Reward selectivity in actor and observer conditions following outcome, data pooled from all sessions in both monkeys. Lower left: scatter plot of actor (x axis) and observer (y axis) reward selectivity, computed as the area under the curve of receiver operating characteristic (ROC) analysis based on spike count in the first 600 ms following outcome feedback. Straight line is the total regression performed on the selectivity values. Black dots are those neurons with significant selectivity in either condition. Green and magenta squares are example neurons from either animal. Top left: histogram of reward selectivity in the actor condition. Black bars correspond to neurons with significant selectivity (890/1628; permutation test, 1000 times, P<0.05). Right: same histogram for the observer condition (significant neurons: 478/1628). Top right: number of neurons that are reward selective in the actor condition (890), observer condition (478), and both (299). (E) Same as (D) for the 600 ms before animals make choice contact in the next trial. Light green and magenta squares indicate the previous selectivity of example neurons in the outcome phase; arrows point to the new selectivity values in the choice phase. (F) Firing rate histograms of example neurons as shown in (D) and (E). Top: Neuron from M1. First vertical dashed line is aligned to outcome, second to choice contact time. Green lines are reward conditions for outcome phase, and previous trial reward for choice phase. Red lines are unrewarded trials. Solid lines are actor conditions for the outcome phase, and previous trial actor conditions for the choice phase. Dashed lines are observer conditions. The gray bars correspond to the 600 ms windows following outcome or before choice used for analysis in (D) and (E). Bottom: same as top for example neuron from M2. (G) Histogram of accumulation selectivity defined as the area under the curve of ROC analysis based on spike count in the first 600 ms following outcome feedback, performed on the rate difference between 1NR vs 2NR, separated into 1NR-Actor, 2NR-Actor (AA), AO, OA, OO conditions. Neurons with significant actor and/or observer reward selectivity with the same sign are included in this analysis. Black bars correspond to neurons with significant selectivity in any of the four conditions (111/630; permutation test, 1000 times, P<0.05). (H) Firing rate histogram of example neurons with significant accumulation selectivity. Top: example neuron from M1. Dashed line aligned to outcome. Green: 1R trials. Light red: 1NR trials. Dark red: 2NR trials. Gray bar corresponds to the 600 ms window used for analysis in (G) and (I). Bottom: same as top for example neuron from M2. (I) Relative change in z-scored firing rate in the 600 ms window following outcome feedback between 1R and 1NR or 2NR. Neurons with significant accumulation selectivity are included in this analysis. Solid line corresponds to actor conditions in 1NR and 2NR, dashed line corresponds to observer conditions. For neurons with decreased firing rate in 1NR vs. 1R, the rate change is multiplied by −1 for both 1NR and 2NR. Error bars indicate 95% confidence interval from bootstrap, N=1000.

A common observation was that single neurons could not be straightforwardly classified in terms of their selectivities. Instead, most task-modulated neurons were sensitive to multiple variables and their sensitivity could change throughout the trial (Figure 3A-C). For example, using receiver operating characteristic (ROC) analysis, we found mixed selectivity to trial outcome for the actor and observer (Figure 3D,E,S6A-D) with a large proportion sensitive to actor outcome (bootstrap test, p<0.05 in 890/1628=54.7% of all neurons in both animals), a smaller proportion to observer outcome (bootstrap test, p<0.05 in 478/1628=29.4% of all neurons in both animals), and a sizeable overlap between the two (299/1069=28.0% of outcome selective neurons). Moreover, this sensitivity changed throughout the trial, as evident from single neurons (Figure 3F) and across the population (Figure 3D-E, regression slopes). Notably, the alignment for outcome encoding between the actor and observer increased in the choice phase compared to the outcome phase (Figure 3D-E; regression slope in choice: 0.46±0.03; outcome: 0.13±0.01; variance explained in choice: 32%; outcome: 6%). This result is consistent with a gradual integration of distinct actor- and observer-dependent responses into an identity-agnostic outcome representation.

Since the animals integrated switch evidence across trials, we further analyzed single neurons for evidence of outcome integration beyond the current trial. A neuron can support this integration if firing rate modulations in the actor and observer trials are in the same direction. Otherwise, evidence from actor and observer trials could negate one another. Therefore, we restricted our analysis to 630 neurons with same-sign outcome selectivity for actor and observer trials (Figure 3D-E, 1^st^ and ^3rd^ quadrants) and used ROC analysis to quantify the difference between firing rates in the 1NR and 2NR conditions (Figure 3G, S6E,F). Across this population, 17.6% (111/630) were more strongly modulated for 2NR compared to 1NR trials. This effect was also evident in the average firing rates of individual neurons (Figure 3H).

Previous work has shown that ACC neurons integrate cross-trial evidence in single-player tasks that involve only experiential evidence (*3*, *10*, *23*). As such, it is critical to subdivide 2NR trials and distinguish between Actor-Actor trials that involve only experiential evidence and the other three conditions that have at least one observer trial (Actor-Observer, Observer-Actor, and Observer-Observer). Doing so, we found that 91.0% (101/111) of neurons that featured evidence accumulation were sensitive to observer trials (Figure 3G, inset).

A notable feature of behavior was the asymmetry in evidence accumulation: unrewarded trials, matched in every other aspect, were more strongly weighed in actor trials compared to observer trials. Accordingly, we asked whether firing rate changes across neurons featuring evidence accumulation were stronger for actor trials compared to observer trials. Results provided strong evidence that firing rate modulations in the 1NR and 2NR trials compared to rewarded trials were stronger in the Actor condition (Figure 3I, S6H-I, mean z-scored rate change in actor condition: 0.36 in 1NR, 0.42 in 2NR; observer condition: 0.12 in 1NR, 0.20 in 2NR; p<0.001 between Actor and Observer conditions, paired t-tests).

### The neural geometry of multi-agent evidence accumulation

A general observation in our single-neuron analyses was that activity patterns were remarkably heterogeneous. While single neurons encoded a wide range of task variables, this sensitivity was typically mixed, changing both during the trial (Choice versus Collect versus Outcome phases) and as a function of trial type (e.g., Actor versus Observer). This property, which is common across frontal cortical neurons (*46*), motivated further neural analysis at the population level, which can offer complementary computational insights (*47–49*).

#### A stable dimension encoding switch belief (Figure 4A-D)

First, we asked whether there exists a dimension within the state space that consistently encodes switch behavior. Using targeted dimensionality reduction (*47*), we identified a dimension along which activity increased monotonically across trials leading to a behavioral switch (Figure 4A). Using cross-validation, we verified that this dimension predicted switch behavior accurately (Figure 4B; Figure S7A-B for individual animals). Importantly, this effect was specific to trials preceding switches and was not due to a trial-order effect (Figure 4B, dashed). We also confirmed the link between the encoding dimension and switch behavior by verifying that large and small projections of neural activity on the encoding dimension corresponded to high and low values of P(switch), respectively (Figure 4C).

**Figure 4.**
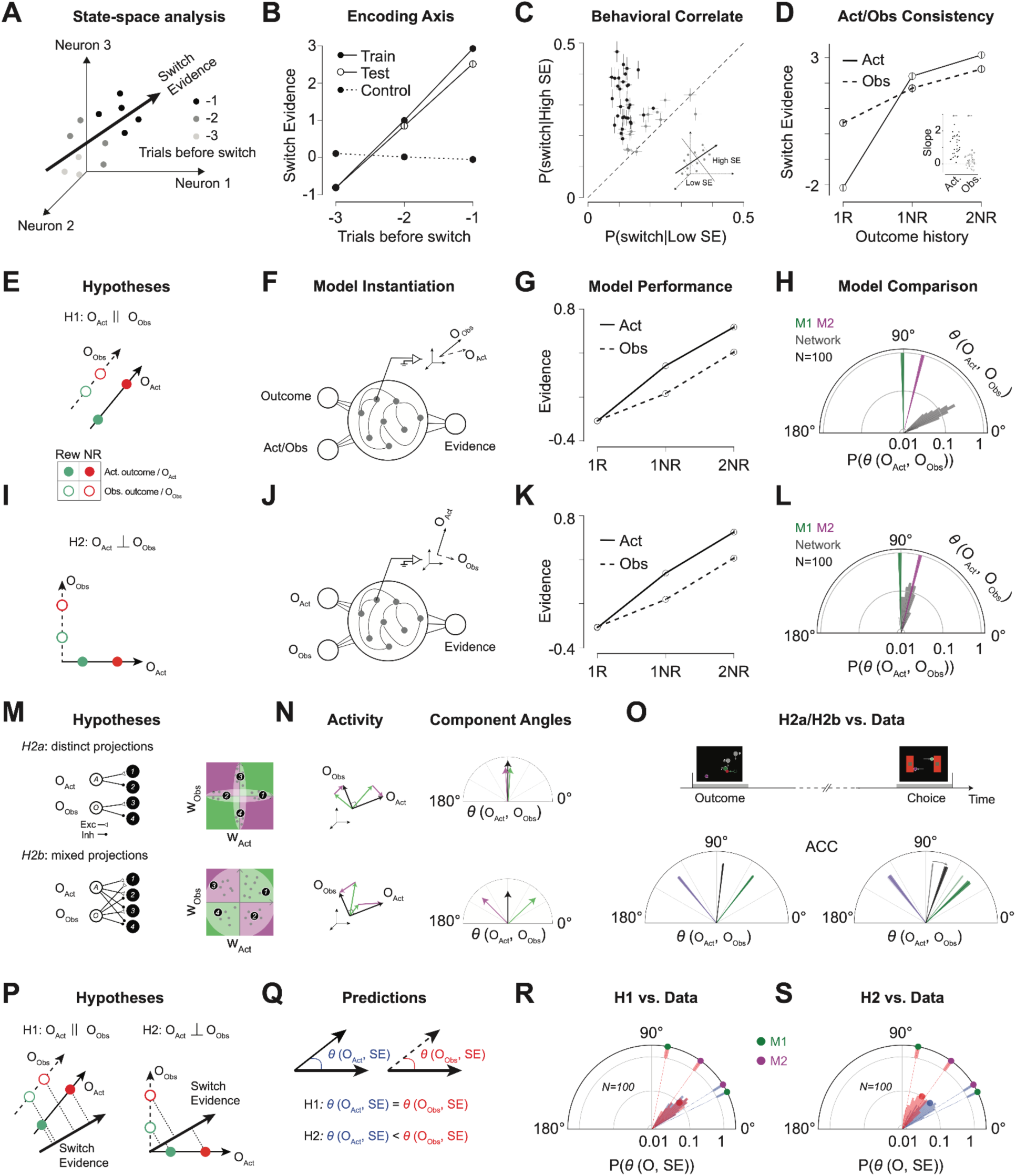
Population geometry of multi-agent evidence integration. (A) A neural switch evidence dimension was computed by regression on the number of trials before switch. (B) Projection on the identified switch evidence dimension relative to number of trials before switch for the data used to compute the dimension (solid disks with solid line), held out test trials (open circles with solid line), and a null control which is selected by treating every three trials as 3,2,1 trials before switch, irrespective of actual switch trials. Data pooled from both monkeys across sessions. (C) Proportion of switch trials conditioned on a low (x axis) or high (y axis) projection on the switch evidence dimension (SE) for each session. Error bars: SEM. Black dots represent sessions with significant differences between low and high projection groups (N=31/43 sessions, rank sum test, P<0.05). Inset: illustration of the procedure. Trials are sorted by the projection on switch evidence dimension and labeled as high or low SE based on the projection value greater or less than the median for each session. (D) Projection on switch evidence dimension by behavioral condition. Solid line: actor conditions. Dashed line: observer conditions. Inset: slope of regression line for projection on SE over outcome history in actor and observer conditions. Data pooled from both monkeys across sessions. (E) Parallel hypothesis on the organization of neural activity. Data pooled from both monkeys across sessions. Each dot represents the average neural activity for one condition, and horizontal lines represent the dimension for actor (solid line) and observer (dashed line) outcomes. (F) Recurrent neural network instantiating the parallel hypothesis. Inputs represent the identity-agnostic outcome and indicator variable for actor or observer conditions. A trainable RNN in the center learns to generate an output that is the integrated evidence. Inset: in-silico physiology was performed to determine outcome and evidence dimensions in artificial neural space. (G) Performance of trained RNNs in (F) on the multi-agent evidence integration task. Y axis is the mean-subtracted model output averaged over 100 model instances. Solid (dashed) line represents output when the indicator variable is set to actor (observer) condition. Error bar: 95% confidence interval. (H) Angle between actor and observer outcome dimensions (θ(O_Act_, O_Obs_)). Gray: θ(O_Act_, O_Obs_) computed from network activation at stimulus onset, repeated for 100 networks, with probability on the radial axis, plotted in 3 degree bins. Green (magenta): θ (O_Act_, O_Obs_) computed from ACC neural activity in M1 (M2) using randomly selected half of trials, repeated 100 times, width indicates standard deviation. (I) The orthogonal hypothesis on the organization of neural activity. Actor and observer outcome dimensions are orthogonal. (J) Recurrent neural network instantiating the orthogonal hypothesis. Same as (F) except the inputs are identity-specific outcomes (one for actor and one for observer condition). (K) Same as (G) for the output from the orthogonal network in (J). (L) Same as (H) except gray bars represent θ (O_Act_, O_Obs_) computed from network activation in the orthogonal network. (M) Hypotheses for the network architecture responsible for orthogonal neural activity. Left: schematic of effective synaptic connections between input populations representing the actor and observer outcome (A and O) and ACC neural population (1–4). Open triangles represent excitatory input and solid circles represent inhibitory input. Right: Each dot represents input weights for one neuron, where the horizontal axis is the weight for the actor outcome, and the vertical axis is the weight for the observer outcome. 1-4: the position of neurons from the schematic on the left in weight space. (N) Left: actor and observer outcome dimensions (black) decomposed into aligned (green) and anti-aligned (magenta) subspaces. Right: Angle between actor and observer outcome dimensions (black), and between aligned (green) or anti-aligned (magenta) subspaces. (O) Decomposition of angles as in (N) for actor and observer outcome dimensions in ACC, computed from the 600ms after outcome (left), and the 600ms before choice (right). Arrow in the right plot indicates change from outcome phase (light shade) to choice phase. Data pooled from both monkeys across sessions. (P) In the parallel hypothesis (H1, left), the switch evidence dimension is parallel to the outcome dimensions. In the orthogonal hypothesis (H2, right), the switch evidence dimension lies between the actor and observer outcome dimensions. Evidence integration occurs by projection from outcome to evidence dimension in both hypotheses. (Q) The angle between the actor/observer outcome dimension and switch evidence dimension in H1 and H2. (R) Angle between actor/observer outcome dimensions and the switch evidence dimension (θ(O_Act_, SE), θ(O_Obs_, SE)). Center blue angular distribution shows values of θ(O_Act_, SE) for 100 H1-instantiating RNNs (F) at stimulus onset. The red shows the same for θ(O_Obs_, SE). Overlaying blue/red circles show the corresponding mean values. Blue dashed line shows average θ(O_Act_, SE) computed from ACC data, separately for the two animals (M1: green; M2: magenta). The shaded blue near the outer circle shows the corresponding std values computed from randomly selected halves of trials, repeated 100 times. Red dashed line and outer circle shows the same for θ(O_Obs_, SE). (S) Same as (R) for H2, with network data from (J).

Since animals’ behavior was sensitive to evidence from both Actor and Observer trials, we next examined projections onto the encoding dimension separately for those two trial types. The encoding dimension carried information for both trial types, with larger projections in the 2NR compared to 1NR trials (paired t-test, p<0.001 in both Actor and Observer conditions, Figure 4D; Figure S7 E-F for individual animals). This dependence on outcome was robust as evidenced by the positive regression slopes relating the projections to the number of unrewarded trials within individual sessions (Figure 4D, inset). Finally, consistent with the asymmetry of evidence accumulation between Actor and Observer trials, the regression slopes were significantly larger for Actor compared to Observer trials (p<0.001, paired t-test, Figure 4D, inset).

#### Identity and outcome inputs to ACC (Figure 4E-L)

We analyzed the neural state space to investigate the arrangement of inputs to ACC that carry information about identity and outcome. We considered two hypotheses. H1 posits that ACC receives the experienced and observed outcomes through a common identity-agnostic input leading to parallel dimensions of representation for Actor and Observer outcomes (Figure 4E). In this scenario, the asymmetry between Actor and Observer has to be accounted for by another input signaling identity (Actor versus Observer). H2, in contrast, posits independent inputs for experienced and observed outcomes leading to orthogonal dimensions of representation (Figure 4I).

To test these hypotheses, we compared the geometry of population activity in ACC to that of recurrent neural network (RNN) models instantiating those hypotheses. For H1, the RNN receives one input supplying identity-agnostic outcome information and another signaling the identity (Figure 4F). For H2, the RNN receives separate inputs for experienced and observed outcomes (Figure 4J). We trained 100 randomly initialized RNNs for each hypothesis on a simplified version of the multi-agent integration task, requiring the models to integrate experienced and observed evidence and replicate the Actor/Observer asymmetry (see Methods).

We confirmed that all RNNs integrated evidence and reproduced the asymmetry in behavior (Figure 4G,K, Figure S8). We then compared the representational geometry of outcome in the RNNs to that of the ACC. Using targeted dimensionality reduction, we identified the Actor and Observer encoding dimensions across datasets and evaluated H1 and H2 by comparing the angle between these two dimensions in the models and ACC.

We compared RNNs instantiating H1 and H2. The angle associated with Actor and Observer dimensions were relatively small for H1, closer to parallel (Figure 4H; 30.28±6.74 degrees, mean±SD, N=100 networks), and larger and closer to orthogonal for H2 (Figure 4L; 79.42±7.00 degrees, mean±SD, N=100 networks, Figure 4L). The corresponding angles in ACC data were also large (90.83±1.35 degrees in M1, 75.97±1.28 degrees in M2, mean±SD, N=100 splits per animal) and fell within the distribution predicted by H2 (Figure 4H,L). These findings suggest that ACC likely receives Actor and Observer outcome information via independent input pathways.

#### Input projection patterns onto ACC (Figure 4M-O, S9A-C)

We analyzed population responses in ACC to dissect the organization of input projections onto ACC. Our previous analysis indicated that ACC receives independent inputs associated with actor and observer outcomes. This orthogonality is consistent with two hypotheses, denoted H2a and H2b. H2a posits that the actor and observer inputs project to disjoint ACC subpopulations (Figure 4M, S9A). This organization is consistent with the input orthogonality because disjoint populations are inherently orthogonal. H2b, in contrast, posits mixed projections to the same population in ACC (Figure 4M, S9A), which could also result in orthogonality.

Many neurons were sensitive to both actor and observer outcomes (Figure 3D, S6A-B). This result provides evidence for some level of mixed projection. To distinguish between H2a and H2b more definitively, we divided the neurons into two groups. The first group, which we refer to as aligned, include neurons whose firing rates move in the same direction for actor and observer conditions, either positively or negatively (Figure 4M, S9A green). The second group, which we refer to as anti-aligned are the neurons that encode outcome with opposite signs for actor and observer (Figure 4M, S9A magenta).

Critically, H2a and H2b make distinct predictions about the geometry of activity within the subspaces formed by these groups. In H2a, where projections are disjoint, the angles between actor and observer outcome representations remain orthogonal for both aligned and anti-aligned neurons (Figure 4N, S9B-C). In contrast, H2b predicts a divergence in the angle relationships: for aligned neurons, the encoding dimensions are more aligned, resulting in acute angles, whereas for anti-aligned neurons, the encoding dimensions are oppositely oriented, leading to obtuse angles (Figure 4N, S9B-C).

In ACC, at the time of outcome, the decomposed angle between actor and observer dimensions had large positive and negative components, consistent with H2b – not H2a (angle at outcome: 83.46° ± 0.92°, Figure 4O left; Figure S7G-H for individual animals). As the task proceeds to time of choice for the next trial, the two vectors become more aligned (angle at choice: 69.94° ± 1.98°, Figure 4O right), consistent with the increased correlation of single neuron selectivities (Figure 3E). Together these results suggest that the actor and observer outcome inputs drive ACC through mixed projection patterns.

#### Geometry of evidence integration (Figure 4P-S)

For effective integration, the switch evidence dimension should be outside the null space of both actor/observer outcome encoding dimensions. This predicts that the angle between the actor outcome and switch evidence, *θ(O_Act_,SE)*, as well as the angle between the observer outcome and switch evidence, *θ(O_Obs_,SE)* must be less than 90 deg. However, the relative geometry of these dimensions differ under the two hypotheses (Figure 4P). Under H1, because the outcome encoding dimensions are parallel, *θ(O_Act_,SE)* and *θ(O_Obs_,SE)* must be the same (Figure 4P). In contrast, under H2, because the outcome encoding dimensions are orthogonal, *θ(O_Act_,SE)* and *θ(O_Obs_,SE)* can assume different values (Figure 4Q).

We applied targeted dimensionality reduction to determine the actor/observer outcomes dimensions as well as the switch evidence dimensions, and used those dimensions to measure *θ(O_Act_,SE)* and *θ(O_Obs_,SE)* in both the models and ACC. Comparing the results, we found that the representational geometry in ACC was better captured by H2 (Figure 4R,S). In both ACC and H2-instantiating RNNs, *θ(O_Act_,SE)* was smaller than *θ(O_Obs_,SE)* (RNN_H2_: 43.15±10.22 vs. 59.11±10.89 deg., M1: 26.25±0.88 vs. 79.05±1.42 deg., M2: 31.96±0.94 vs. 54.40±1.35 deg.). This difference was not driven by an unequal number of Actor and Observer switch trials; results were qualitatively unchanged when we controlled for trial count in the two conditions. (M1: 35.19±1.89 vs. 75.29±2.49 deg., M2: 33.22±1.43 vs. 53.50±1.67 deg.). In contrast, the two angles in the H1-instantiating RNNs had similar magnitudes (44.03±7.35 vs. 43.48±7.80 deg.).

The fact that *θ(O_Act_,SE)* was smaller than *θ(O_Obs_,SE)* is noteworthy in light of the behavioral asymmetry in experiential and observational evidence integration. To be integrated, actor and observer outcomes must be mapped onto a common switch evidence dimension. In H1, the two angles are nearly the same, and therefore, the only constraint is that the switch evidence dimension must be outside the null space of the actor/observer outcome. In contrast, in H2 where the angle can differ, the relative geometry of the switch evidence dimension is consequential. If integration occurs by linear projection of activity along the outcome dimension onto the switch evidence dimension, then a smaller angle between the actor outcome and switch evidence dimensions would enable the same strength of evidence collected on actor trials to produce a higher increment in cumulative evidence (Figure 4P). Therefore, the smaller angle associated with the actor outcome representation in ACC provides a potential mechanistic explanation for the more effective integration of evidence in the actor trials.

So far, we configured all RNNs such that their output weights were fixed. To test the effect of this assumption on our findings, we performed control analyses on RNNs built with learnable output weights (Figure S10B,D). The geometry of evidence integration in these readout-learnable networks was different from both the readout-fixed networks and the ACC. Specifically, they exhibited relatively higher alignment between actor and observer outcome dimensions and outcome and evidence dimensions (Figure S10B,D). These results suggest ACC internal dynamics and not its downstream projections are responsible for evidence integration.

## Discussion

Our work brings together two important yet traditionally distinct areas of research concerning the role of ACC in cognition. One important function of ACC is to monitor and integrate one’s experience over time to inform strategic decision-making (*4*, *11*, *17*). This function supports a wide range of mental computations including explore-exploit trade-offs, cost-benefit analysis, conflict monitoring, and causal inference (*1*, *3*, *9*, *10*, *19*, *23*, *25*). Another function ascribed to ACC is sensitivity to observed reward and punishment enabling vicarious learning (*35*, *36*, *39*– *42*). Our work offers a unifying perspective wherein ACC plays a general role in integrating experiential and observational outcomes over flexible timescales to update belief about environmental states. The confluence of these two research directions brings to focus several important questions.

First, what anatomical substrates and circuit motifs enable the integration of information about self- and other-experience? We found that ACC encodes all three key computational variables needed for integration: actor outcome, observer outcome, and integrated belief. A comparison of population activity between models and ACC provided evidence that actor and observer outcomes were associated with activity patterns in orthogonal subspaces. This finding suggests that ACC computes beliefs about the state of the environment by integrating outcome information from distinct identity-dependent input streams.

Analysis of the geometry of neural representation is often used to infer computational algorithms. For example, subspace orthogonality is thought to prevent interference and maximize robustness (*47*, *50–52*), and factorized representations are thought to facilitate structural generalization (*49*, *53–57*). In our work, we augmented this analysis with single-neuron tuning properties to dissect the organization of input projections onto ACC. Specifically, we analyzed the representational geometry separately for aligned and anti-aligned actor and observer outcome encoding neurons. Results indicated that ACC did not rely on disjoint subpopulations for actor and observer information. Instead, actor/observer information was supplied via overlapping projections. We do not know the constraints that determine the organization of these projections. However, in our experiment, this mixing may facilitate the integration process. With disjoint subpopulations, the integration would have to be augmented by a gating mechanism to select the subpopulation that has to be integrated on each trial. In contrast, the mixed representation provides a single subpopulation of outcome-aligned neurons that can be used for integration in all trials.

Our analyses, both at the single-cell and population levels, indicated that ACC coding properties changed throughout the trial. One notable feature of these dynamics was the reduction of the angle between the population vectors encoding the actor and observer outcomes, from the outcome phase to the choice phase. This finding is reminiscent of prior work in the lateral intraparietal area showing that generically high-dimensional firing-rate vectors rapidly decay to a single dimension during the process of decision-making (*58*). However, in our work, this process unfolded in the presence of multiple inputs (two agents) and long timescales (across trials), which pose important constraints on models of evidence accumulation in ACC (*59*, *60*).

Second, does the brain process experiential and observational information similarly? In our two-player game, although humans and monkeys integrated experiential and observational outcomes, they learned less from observations. Discounting observational evidence has been reported previously (*43–45*). However, several aspects of our study reinforce the view that there is a fundamental asymmetry between learning from experience and observation. By interleaving actor and observer trials while collecting each player’s choice on every trial, we could track both players’ evolving beliefs with precision. This design choice as well as our analysis of congruent and unrewarded trials enabled us to rule out various confounds that could lead to this asymmetry. Finally, we found this asymmetry to be stronger in monkeys, possibly because monkeys – not humans – received juice reward, which could accentuate the difference between experience and observation. The difference between humans and monkeys may also stem from superior social cognition in humans enabling more effective evaluation and integration of observations.

We identified a neural correlate of this asymmetry in the ACC, where signals encoding actor outcomes were more closely aligned with cumulative switch evidence than those encoding observer outcomes. Validating the functional relevance of this finding will require precise patterned activations of subpopulations of neurons in ACC (*61–63*). Additionally, characterizing the behavioral contingencies and neural constraints that give rise to this asymmetry remains an important direction for future research.

In sum, our work establishes the basic mechanisms of multi-agent evidence integration and offers a starting point for addressing exciting and unresolved questions about social learning. Extensions of our work can be used to study the mechanisms through which cognitive factors such as belief about the partner’s skill level, their prior knowledge about task contingencies, and their social rank influence observational learning.

## Materials and Methods

We collected behavioral data from humans, and behavioral and neurophysiological data from rhesus macaque monkeys (Macaca mulatta). Experimental procedures for humans were approved by the Committee on the Use of Humans as Experimental Subjects at the Massachusetts Institute of Technology. Experimental procedures for animals conformed to the National Institutes of Health guidelines and were approved by the Committee of Animal Care at the Massachusetts Institute of Technology.

### Experimental procedures for non-human primates

Two monkeys (M1, female, 6 Kg, aged 6; M2: male, 11 Kg, aged 11) were seated comfortably in two adjacent primate chairs in a dark quiet enclosure at a distance of 40 inches. Animals were head-restained, facing forward, and unable to see one another. Stimuli were presented on two side-by-side display monitors (Acer R240HY) 40 inches apart (center to center), at a normal distance of 19 inches from the animals’ eyes. The monitors displayed identical stimuli throughout experiments. Each animal could manipulate a joystick (Logitech Extreme 3D Pro) placed at a distance adjusted for each animal’s reach (∼6 inches) in front of their chair. Joysticks were physically constrained to left/right movements only. The joystick digital output (1–1024) was thresholded to three states: left movement (1–463), no movement (464–560), and right movement (561–1024). Eye movements were sampled at 1kHz using infrared cameras (Eyelink 1000, SR Research). The MWorks software package (http://mworks-project.org) and MOOG library (*64*) (https://jazlab.github.io/moog.github.io/) were used to generate visual stimuli and to enforce behavioral contingencies. A photodiode was used to sync electronic events with stimulus presentations.

Neural recordings were made from the anterior cingulate cortex (ACC) with 64-channel linear probes with 50-µm inter-electrode spacing (V-probe, Plexon Inc.) inserted through a rectangular recording chamber. Extracellular signals were bandpass filtered (300 Hz to 6 kHz) and digitized (sampling rate: 30 kHz) using two 32-channel headstages (Intan Technologies), and collected using OpenEphys software (http://www.open-ephys.org). Spike sorting and curation were carried out using Kilosort 3 (https://github.com/MouseLand/Kilosort) and phy (https://github.com/cortex-lab/phy). Recording sites and number of sessions/trials are reported in Table S1. Data analysis was performed using custom Python code.

### Experimental procedures for humans

We recruited a total of 14 participants. All participants gave informed consent, were naive to the purpose of the study, and had normal or corrected-to-normal vision. Participants were asked to play the single-player version of the task first, and after gaining familiarity were invited to play the two-player version. One participant did not learn the task after three single-player sessions (set to be an exclusion criterion before data collection begins). Three participants withdrew voluntarily after finishing the single-player sessions. The remaining 10 participants (6 males and 4 females, aged 18-65) completed the two-player sessions. These participants were divided into 5 fixed pairs (1 female-female, 2 male-male, and 2 male-female). Each session lasted ∼60 min. Participants were paid a fixed amount at the end of each session. Those who finished all single and two-player experiments were paid an additional 50% of all their earnings as a bonus.

In each session, participants sat in adjacent dark enclosures, separated by an opaque curtain to prevent visual contact. Each enclosure had identical equipment (monitor, keyboard, joystick connected to a Mac mini). Participants knew their monitors displayed the same visuals, though they couldn’t see the other screen. In the single-player experiment, only one setup was used. Like monkey experiments, tasks, stimuli, and behavioral contingencies were controlled by MWorks and MOOG software packages.

### Behavioral task

We devised a two-player trial-based game. Each trial consists of two hierarchically organized phases. Phase 1 begins with the presentation of the two arenas on the two sides of the monitor and two disks stacked on top of one another mid-way between the two arenas representing the two players (‘avatars’). Each arena is a red rectangle (8.2 x 16.4cm) with a central gray square (width: 4.1cm) placed 3 cm to the left or right of the vertical midline. Each avatar appears as a yellow disk (diameter: 2.4cm) placed 3.85 cm above or below the horizontal midline. After 750 ms, the two yellow avatars are randomly assigned to the two players by a change of color, purple for player 1, and green for player 2. At this time, players must choose either the left or right arena by moving their avatar in the direction of their preferred arena. Once an avatar contacts an arena, the joystick control for that avatar is temporarily relinquished. After both players choose their preferred arena an additional 750 ms interphase-interval delay is imposed and then the task moves to the second phase. In phase two, players are randomly assigned to be the actor and observer, the red rectangles disappear, and the two avatars are displaced from the edge to the bottom interior of the corresponding arenas. The actor is placed 10.1 cm away from the vertical midline and 6 cm below the horizontal midline. The observer is placed 10.1 cm away from the vertical midline and 12 cm below the horizontal midline. Immediately afterward, the actor starts playing the token capture game on its monitor, a copy of which is shown on the observer’s monitor. The actor must use the joystick to move the avatar left or right to capture 15 falling tokens (gray circular disks of diameter: 6.72 cm). The tokens drop sequentially every 1/3 s, starting 16.8 cm above the center and moving directly downward at the constant speed of 50.4 cm/s. The horizontal positions of tokens are sampled from a Gaussian Process (GP) centered at the initial position of the actor’s avatar with a squared exponential kernel. To vary task difficulty across trials, the standard deviation of the GP was sampled from a discrete uniform distribution ([2.52, 5.04, 7.56] cm).

Each trial ends in either a win or a lose state. To maximize wins, the actor must select the correct arena and collect as many tokens as possible. The correct arena (the one associated with a non-zero win probability) switches covertly in a blocked fashion (see below). In either arena, captured tokens turn green, while missed tokens disappear without effect. A win is signaled by a change of the color of the avatars and an auditory tone. A trial ends either with a win or after all 15 tokens pass. The observer receives no reward but can monitor trial progression and infer the outcome from the visual and auditory feedback. After the trial, the display remains stationary for 1 second before switching to a uniform black screen for an inter-trial interval of 2.676 ± 0.085 seconds.

Before the two-player game, players completed a single-player version with a single avatar, designated as the actor. All other task aspects remained identical to the two-player version. The task contingencies were identical for humans and monkeys with a few exceptions as follows.

#### Non-human primates

(a) The switching probability is 0 for the first 10 trials, 1/3 for trials 11-24, and 1 for trial 25. (b) In the correct arena, captured tokens may trigger a win with a 15% probability. (c) A win triggered a juice drop for the actor. (d) Animals first played the single-player version until their performance stabilized. The single-player data reported are from sessions with stable performance. (e) They were then introduced to the two-player version without electrophysiology recordings until performance stabilized. Reported behavioral and neural data are from subsequent sessions with simultaneous behavioral and physiological recordings.

#### Humans

(a) The switching probability is 0 for the first 10 trials, 1/3 for subsequent trials. (b) In the correct arena, captured tokens may trigger a win with a 10% probability. (c) Before the start of the first session, participants read verbal instructions about the game but were not informed of the reward probabilities or state-switching statistics. (d) Participants played the single-player version for 5 or 6 sessions (see below). Single-player sessions started with 50 practice trials where the correct and incorrect arenas were cued (green and yellow, respectively). After an optional short break, participants completed 600 trials of the one-player game with 2-minute breaks every 200 trials. Reported single-player data are from sessions 1-5 and do not include the practice trials. (e) Initially, participants were scheduled for five single-player sessions. During data collection, a confidence-reporting step was added, requiring participants to use a keyboard to report their confidence on every trial (1: “not confident at all” to 4: “fully confident”) after selecting the arena. Nine participants who completed five sessions in the original task played a sixth session for confidence reporting, while the tenth participant used the augmented task from the start and did not require a sixth session. (f) Participants received instructions about the two-player game before the first two-player session. Each session included 525 trials with two 2-minute breaks after every 175 trials. Each pair completed 15 two-player sessions, totaling 7875 trials.

### Analysis of behavior

We used players’ choice behavior to quantify the probability of choosing the correct arena from 4 trials before to 10 trials after a block switch (Figure 1H-K).

#### Solipsistic agent

To test if a player was sensitive to observer trials, we compared its performance to the performance predicted if the player were to ignore all observer trials and treat the two-player game as a single-player. We refer to this hypothetical player as *solipsistic*. If we denote a win state at trial *n* by *R_n_*, then the average probability of *R_n_* for a solipsistic actor in the two-player game (*P_2p_*) based on their average performance in the one-player game (*P_1p_*) can be written as follows:

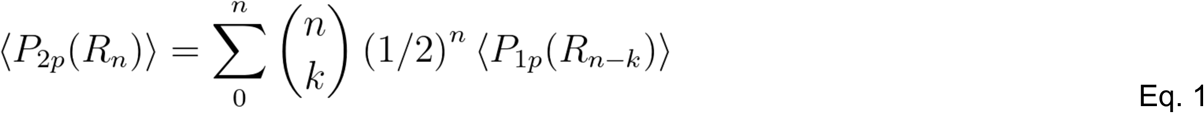

The <.> denotes average probabilities. The sum runs over the Observer trials preceding trial *n* since the last block switch, indexed by *k*. To understand this equation, consider a sequence of *n* trials with *k* Observer and *n-k* Actor trials. The binomial coefficient (*n*-choose-*k*) counts such combinations, while *(½)^n^* gives their probability, with *(½)^k^* for *k* Observer and *(½)^n-k^* for *n-k* Actor trials. The term *P_1p_(R_n-k_)* implements the assumption that the actor disregards Observer trials, behaving as if only *n-k* trials have passed since the last block switch.

We tested the significance of the difference in accuracy between this agent and the participant using t-test on the average P(correct) for positions [0,10].

#### Oracle agent

To estimate an upper performance bound on trial *n*, we simulated an oracle agent who mimicked the player’s choices for trial *1:n-1* but selected the correct choice on trial *n*. Consequently, the oracle chose the correct arena immediately upon an objective switch.

### Difficulty of the collecting tokens

In the second stage, token positions were random, creating varying difficulty levels. To assess whether players attempted to maximize captured tokens, we analyzed performance on unrewarded trials as a function of difficulty (*D*), defined as the sum of absolute distances between successive tokens.

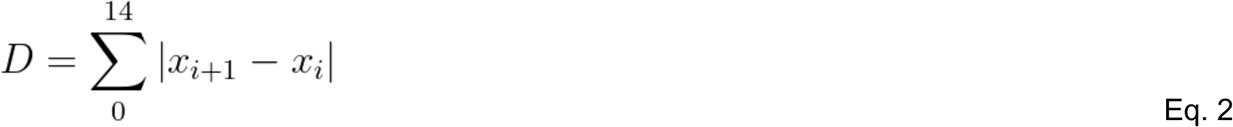

Here, *x_i_* is the horizontal position of the i-th token in screen coordinates. Since each unrewarded trial has 15 tokens, the difficulty is the sum of 14 horizontal displacements.

### Subjective belief about the correct arena and block switches

Since the correct arena and block switches were covert, we developed a method to estimate participants’ subjective beliefs about the arena and trial position within a block. The first rewarded trial of a session was assumed to indicate the correct arena and was assigned position 1 of the first block. The position incremented until the first reward on the opposite arena, which was assumed to signal a block switch, resetting the position to 1. Early, Mid, and Late trials were defined as [1,5], [6,10], and [11, ∞), with bin sizes balanced as closely as possible.

### Regression analysis of switch behavior and confidence

We used logistic regression to quantify the dependence of switch behavior on various factors after unrewarded trials, including the number of consecutive unrewarded trials, the number of tokens captured, and position in the trial.

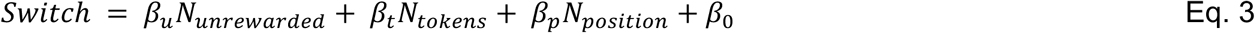

Switch is a binary variable indicating when the player chooses a different side than the actor’s current choice (i.e., the arena for which direct evidence is acquired). *N*_*unrewarded*_ is the number of consecutive unrewarded trials in the same arena. We only included up to 4 consecutive unrewarded trials in this analysis. *N*_*tokens*_ is the number of touched tokens, and *N*_*position*_ is the subjective trial position in the block.

Combining both Actor and Observer trials, we used simple logistic regression (for each monkey) or mixed effects logistic regression (for human participants):

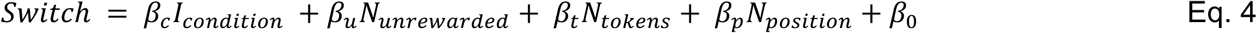

*I*_*condition*_ is a binary indicator variable (0 for Actor, 1 for Observer).

We also computed the contribution from choice conflict for the Actor condition. For this analysis, we included incongruent trials, which were excluded in previous regressions.

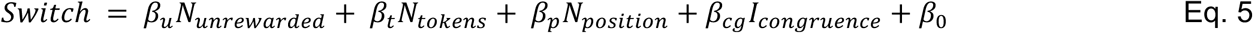

*I*_*congruence*_ is a binary indicator variable (1 for incongruent, 0 for congruent).

For human participants, we additionally performed mixed effects linear regression on their confidence report.

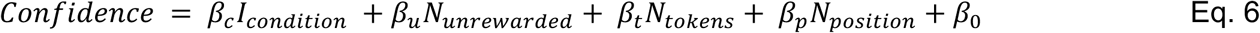

### Analysis of single neurons

We estimated each neuron’s firing rate by averaging spike counts in shifting 100 ms time bins with a 10 ms step size. We analyzed firing rates aligned to different task events including the choice time (i.e., when the avatar contacts an arena) and outcome (i.e., reward time in rewarded trials and end of token collection in unrewarded trials). For visualization, binned firing rates were smoothed using a 3-bin moving average.

### Single-neuron sensitivity to outcome and choice

We measured selectivity to reward outcome in self/other conditions using receiver operating characteristic (ROC) analysis based on spike counts within 600-ms windows, either after the outcome or before the choice. The ROC score, calculated as the area under the performance curve of a binary classifier, classified trials as rewarded or unrewarded—either for the current trial (outcome) or the previous trial (choice). To center selectivity at 0, we subtracted 0.5 from the score, where values of −0.5 and 0.5 indicate perfect separation of reward outcome with lower or higher firing rates for the reward condition, respectively. Significance was assessed via bootstrap (1000 iterations, p<0.05). To compute the correlation of ROC scores between self and other conditions, we performed total least squares regression between self and other selectivities.

### Single-neuron sensitivity to cumulative errors

We computed selectivity for the history of congruent unrewarded trials (1NR vs. 2NR) using ROC analysis for neurons that exhibited significant reward selectivity in either the Actor or Observer condition and maintained the same selectivity sign in both. Binary classifiers were constructed for the four possible consecutive outcomes: 1NR-Actor vs. 2NR-Actor (AA), 1NR-Actor vs. 2NR-Observer (AO), 1NR-Observer vs. 2NR-Actor (OA), and 1NR-Observer vs. 2NR-Observer (OO). For neurons with significant selectivity in any condition, we also calculated the average differences in z-scored firing rates across rewarded Actor trials, 1NR (Actor or Observer), and 2NR (Actor or Observer). Neurons with consistent rate changes (either 1R<1NR<2NR or 1R>1NR>2NR) were classified as encoding cumulative error.

### Population activity for actor/observer outcome and switch evidence

We used targeted dimensionality reduction (TDR) to identify encoding dimensions of actor outcome, observer outcome, and switch evidence (*47*). To do so, we used regression to relate each neuron’s spike count within a 600 ms window following the outcome to different task variables.

For Actor/Observer outcome (analyzed separately), we used the following regression:

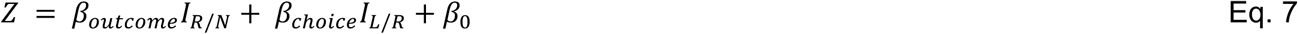

*I*_*R*/*N*_ is an indicator variable for outcome (1 for unrewarded, 0 for rewarded) and *I*_*L*/*R*_ an indicator variable for choice (−1 for left, 1 for right). To reduce estimation noise, only neurons that had more than 5 trials in all conditions were included.

For switch evidence, we used the following regression:

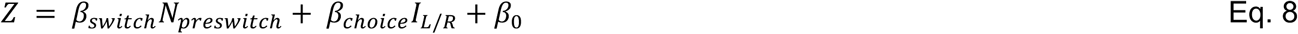

*N*_*preswitch*_ represents the distance in trials from the next switch, with values of −1 for one trial before, −2 for two trials before, and −3 for more than two trials before the switch. *I*_*L*/*R*_ is an indicator variable for choice (−1 for left, 1 for right). To reduce estimation noise, only neurons that had more than 10 trials in all conditions were included.

After solving the regression for all neurons, we arranged the coefficients in a matrix and orthogonalized columns using QR-decomposition, requiring R to have positive diagonal values such that the columns of Q provided are orthogonalized set of coefficients for each variable.

For Actor/Observer (Equation 7), we used the orthogonalized coefficients associated with the outcome (first column) as the Actor/Observer outcome dimension for each condition. For the switch evidence (Equation 8), we used the coefficients associated with *N*_*preswitch*_ (first column) as the switch evidence dimension. For cross-validation of the switch evidence dimension, we randomly selected one trial per condition, computed the evidence dimension from the remaining trials, and projected the held-out trial activity onto this dimension. This process was repeated 100 times, generating 100 cross-validated projections per condition.

### Predicting switch behavior from switch evidence dimension

We projected neural activity onto the switch evidence dimension to predict switch behavior in the next trial. For each recording session, we selected trials with at least two recorded neurons and at least 10 switch trials. Using spike counts from all but one randomly selected trial, we computed the switch evidence dimension and projected activity from all trials onto it, generating a distribution of projection values. The held-out trial was classified as high or low evidence based on whether its projection value was above or below the median of this distribution. This process was repeated 1000 times per session. We then calculated the proportion of switch trials in the high and low evidence groups, considering sessions significant if the low evidence group had a significantly lower proportion of switches, as determined by a rank-sum test.

### Contribution of actor/observer outcome to switch evidence dimension

We computed neural switch evidence in Actor and Observer conditions by randomly selecting one trial from each Actor/Observer × 1R/1NR/2NR condition as a held-out test trial, deriving the evidence dimension from the remaining trials, and projecting the test trial activity onto this dimension. Only accumulation-selective neurons from sessions where neural switch evidence significantly predicted switching behavior were included. This process was repeated 100 times to generate a distribution of projection values for each condition.

### The geometry of actor/observer outcome, and switch evidence

We measured pairwise angles between actor outcome, observer outcome, and switch evidence dimensions using a randomly selected half of the trials and repeated this process 100 times generating a distribution of angles. We also measured the angle between actor and observer separately in two subspaces: the aligned subspace, computed from those neurons whose coefficients in Actor- and Observer-outcome coefficients had the same sign; and the anti-aligned subspace, which had opposite signs.

### Neural network model for multi-agent integration task

We used recurrent neural network (RNN) models with different architectural and optimization constraints to test two hypotheses about how ACC integrates experiential and observational evidence into cumulative switch belief. One hypothesis (H1) posits that ACC receives the experienced and observed evidence through a common identity-agnostic input while another input provides information about the identity (self/other). The other hypothesis (H2) posits that ACC receives independent inputs for experienced and observed outcomes.

#### Architectural constraints

All models have three layers, an input layer providing three distinct inputs, a hidden layer consisting of 200 recurrently connected units, and an output layer for computing the network output.

RNNs instantiating H1 receive one input conveying information about both experienced and observed outcomes and another input for identity (Actor: −1, Observer: 1). RNNs instantiating H2 receive experienced and observational outcomes through separate inputs, with only one of them active in any given trial (the other one is set to zero). In both cases, a third input served as a go cue instructing when the RNN has to generate an output, *Y*, which is a scalar reflecting the cumulative evidence across trials.

For all RNNs, the input carrying outcome information is a sample from a bimodal Gaussian distribution (Equation 9), with one positive and one negative mode centered at 0.5 and −0.5, corresponding to win (rewarded) and lose (unrewarded) states, respectively.

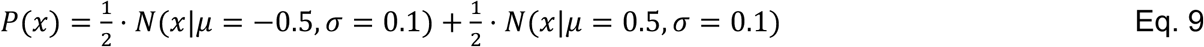

For both H1 and H2, we tested two model variants. In one variant, we assumed the readout weights that drive the output were learnable (H1^learn^, H2^learn^), and in the other, the readout weights were initialized randomly and were not learnable (H1^fix^, H2^fix^). The weights of the input layer were initialized randomly and were not subjected to learning.

#### Optimization constraints

All RNNs were trained to adjust their output, *Y*, according to the following requirements:

(a) *Y* must reset to zero after rewarded trials (positive inputs).
(b) *Y* must integrate outcome input following unrewarded trials (negative inputs) for both Actor and Observer conditions and maintain the integrated value across trials.
(c) To replicate the Actor/Observer behavioral asymmetry, Actor and Observer inputs must be integrated with a gain of 1 and 0.5, respectively.
(d) In the first variant (H1^learn^, H2^learn^), both the recurrent and readout weights were trained. In the second variant (H1^fix^, H2^fix^), training was applied only to the recurrent weights.

Information was presented in a trial-based manner, with each trial having a variable duration sampled uniformly between 10 to 20 timesteps. The input was provided at the first timestep and then set to zero. A go cue input instructed the RNN when to generate an output. This go cue was presented as a linear ramp from 0 to 1 over 5 timesteps, remaining at 1 for an additional 5 timesteps. The onset of the go cue, relative to the trial start, was sampled uniformly from 0 to 5 timesteps. RNNs were required to compute and maintain the output while the go cue is 1.

#### Model dynamics

The activity of hidden units is given by

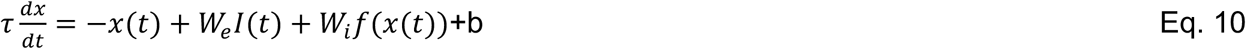

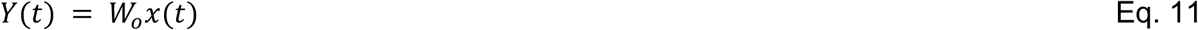

In Equation 10, *τ*=5, *x*(*t*) is the activity of all units, *W*_*e*_ is the embedding weights for inputs, *I*(*t*), *W*_*i*_ is the recurrent weights, *f* is a tanh nonlinear function, and b is a bias term. In Equation 11, *Y*(*t*) is the output and *W*_*o*_ is the readout weights. All parameters were randomly initialized from a normal distribution with zero mean and variance 1/N, where N is the number of parameters for each layer. We trained the networks using gradient descent in batches of 16, by minimizing the MSE loss between output and target (when the go cue is at level 1. We do not constrain network dynamics outside of the reporting window.

We trained 100 models with random initiations per hypothesis, for a total of 400 models, and each model was trained for 100,000 iterations, with 500 timesteps per iteration.

#### Model performance

We computed the performance of trained networks as the average output in the reporting period conditioned on input and trial history. 1R, 1NR, and 2NR were assigned in the same way as in behavioral trials.

#### Model analysis

Similar to ACC, we applied TDR to activity in the hidden layer units to identify encoding dimensions for actor outcome, observer outcome, and switch evidence.

For Actor/Observer outcome (analyzed separately), we used the following regression:

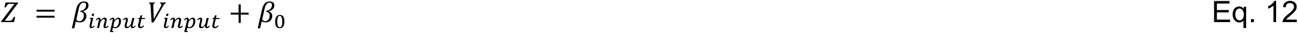

*V*_*input*_ is an indicator variable for outcome (1: unrewarded; 0: rewarded). For switch evidence, we used the following regression:

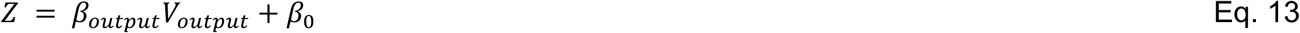

*V*_*output*_ is the output value at the time when the go cue reaches 1 for each trial.

## Acknowledgments

RC was supported by the Simons Foundation and Hock E. Tan and K. Lisa Yang Center for Autism Research. SR was supported by Mathworks Graduate Fellowship and K. Lisa Yang ICoN Center Fellowship. SBY was supported by the Simons Foundation. MJ was supported by the Simons Foundation and the McGovern Institute.

## Author contributions

SR, SBY and MJ designed the task. SBY trained the animals. RC and NV performed animal experiments. SR performed human experiments. RC, SR, and NV analyzed the data. RC, SR, and MJ wrote the manuscript. MJ supervised the project.

## Data availability

The source data used in the manuscript will be made available publicly upon publication.

## Code availability

The custom scripts used for data analysis in this manuscript will be made available publicly upon publication.

