## Supplementary Materials for "Evidence accumulation from experience and observation in the cingulate cortex"

†Equal contribution

**Supplementary Materials**

Figure S1. Eye gaze pattern during token collection

Figure S2. Additional task feature statistics

Figure S3. Effect of observer condition on switching behavior and confidence

Figure S4. Effect of outcome, trial in block, and tokens captured on individual monkeys

Figure S5. Effect of choice congruence on switching behavior and confidence

Figure S6. ACC neurons encode and integrate actor and observer outcome

Figure S7. Population geometry of multi-agent evidence integration

Figure S8. RNN architecture, task, and output

Figure S9. Angle decomposition analysis

Figure S10. Readout-fixed and readout-learnable networks

Table S1. ACC recording location

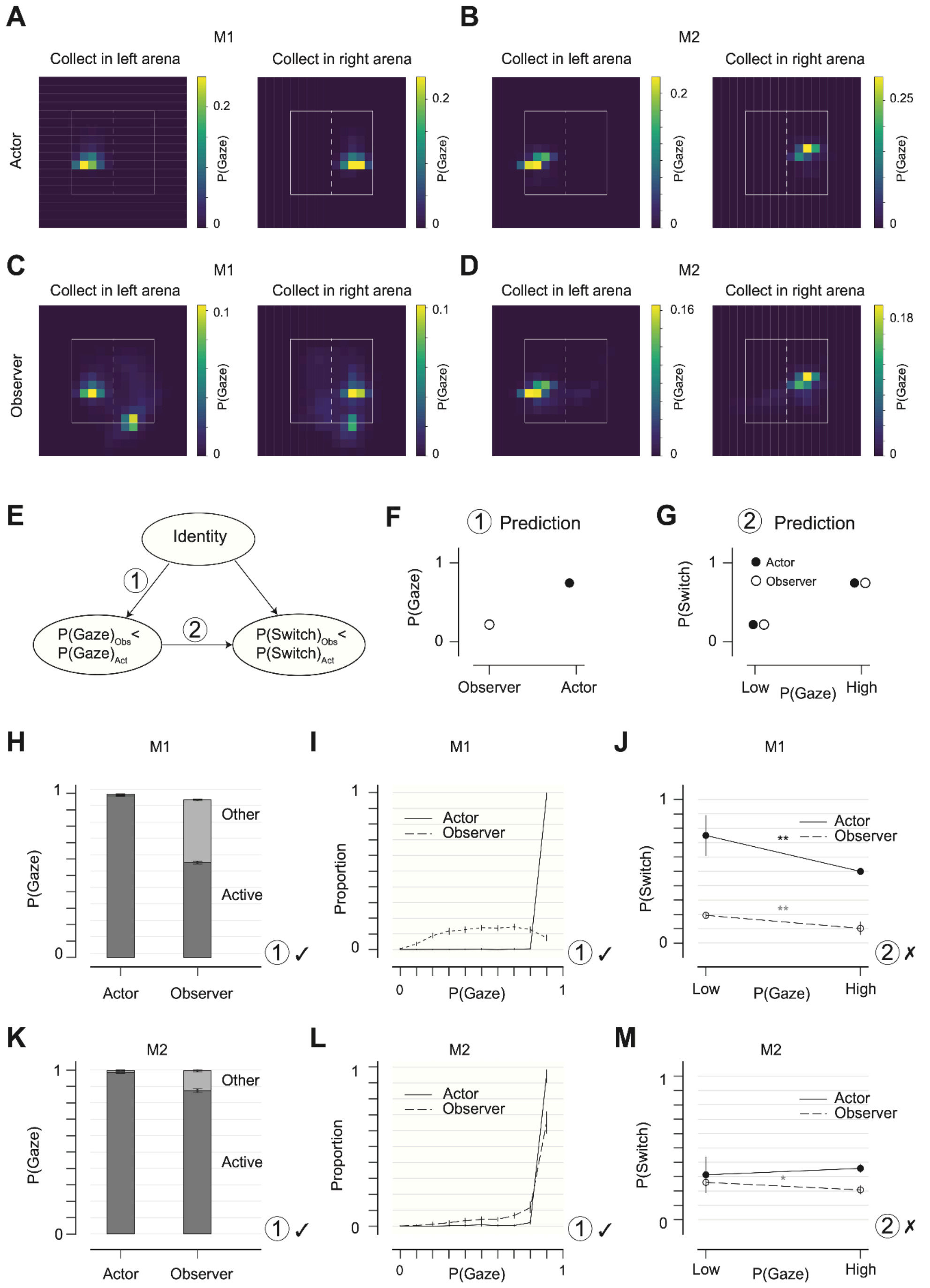


Figure S1. Eye gaze pattern during token collection

(A) Left: distribution of the actor's eye gaze within and near the screen (white square; dashed vertical line indicates center of screen), for congruent left trials. Data from M1 across sessions where eye gaze data was available (26 sessions). Right: same for congruent right trials.

(B) Same as (A) with data from M2 (22 sessions).
(C-D) Same as (A-B) for the observer.
(E) Hypotheses on the causal influence of identity on eye movement and choice behavior. Two hypothetical links are highlighted: (1), Observer condition results in lower P(Gaze); (2), lower P(Gaze) results in lower P(Switch) following unrewarded trials.
(F) Hypothesis (1) predicts lower P(Gaze) in the Observer condition.
(G) Hypothesis (2) predicts equal P(Switch) conditioned on the same P(Gaze).
(H) Proportion of time when eye gaze was within the active/other half of screen during token collection (P(gaze)), calculated as duration of eye gaze in one arena divided by the total duration of collect phase, for all congruent unrewarded trials, data from M1. Error bars indicate 95% confidence interval from bootstrap, N=1000. Hypothesis (1) is supported.
(I) Distribution of P(gaze) on the active side over congruent unrewarded trials, data from M1. Error bars indicate 95% confidence interval from bootstrap, N=1000. Hypothesis (1) is supported.
(J) Probability of switching on the next trial following low (<90%) and high (>90%) P(gaze), data from M1. Error bars indicate 95% confidence interval from bootstrap, N=1000. Stars indicate significant difference between low and high P(gaze) (t-test, **:$P<0.01$, *:$P<0.05$). Hypothesis (2) is not supported.
(K-M) Same as (E-G) with data from M2.

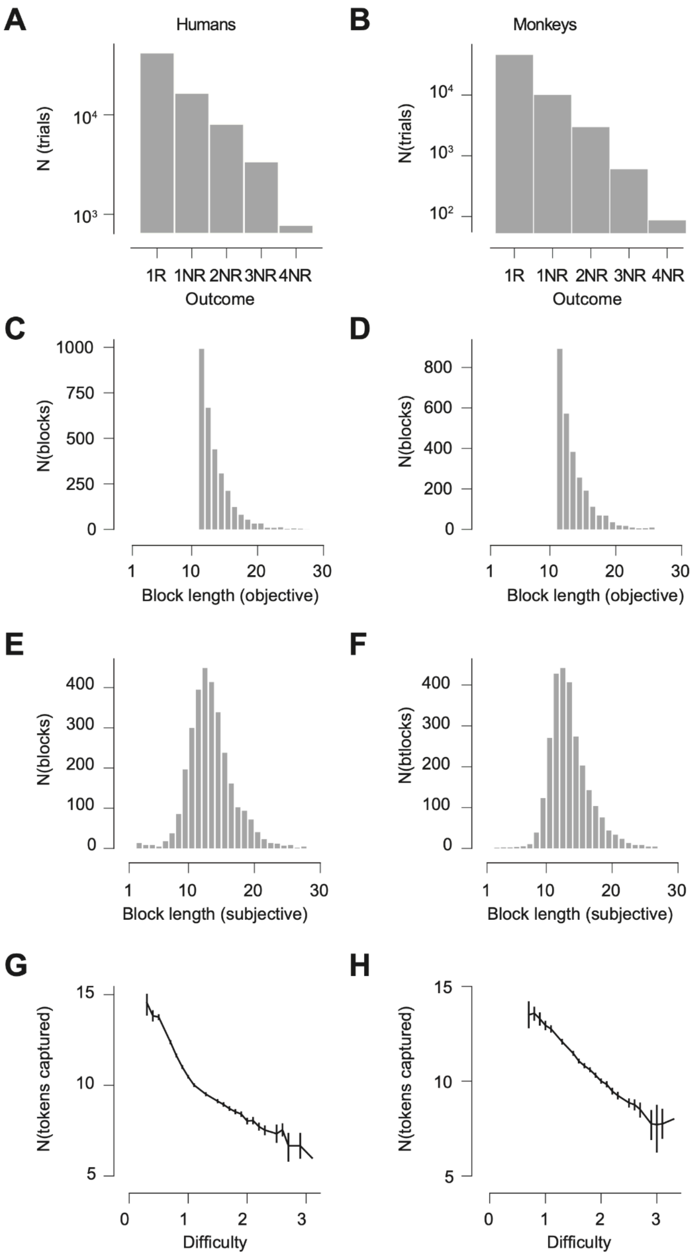


Figure S2. Additional task feature statistics

(A) Number of trials in each outcome group, data from all two-player sessions in human participants. 1R: rewarded trials where both animals made the same choice. nNR: n-th consecutive unrewarded trial where both animals consistently made the same choice as the most recent rewarded trial.

(B) Same as (A) for monkeys.

(C) Distribution of object block length (number of trials) in two-player sessions in human participants. Objective block lengths are computed from covert switches and not cued to participants. One block had a length of 31 trials and is not shown in the plot.
(D) Same as (C) for monkeys.
(E) Distribution of subjective block length in two-player sessions in human participants. Subjective block lengths are computed as trials since the first rewarded trial in the current side (methods). 8 blocks had lengths greater than 30 and is not shown in the plot.
(F) Same as (E) for monkeys.
(G) Number of tokens captured by the actor as a function of difficulty. Difficulty is computed as the cumulative horizontal displacement between successive tokens (methods). Vertical lines indicate 95% confidence interval.
(H) Same as (G) for monkeys.

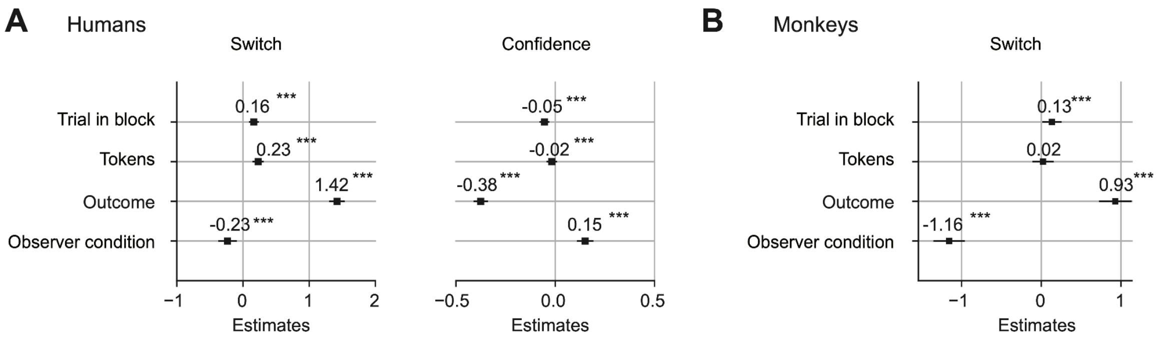


Figure S3. Effect of observer condition on switching behavior and confidence

(A) The effect of condition ('Observer' vs 'Actor') on switching behavior and reported confidence, while controlling for outcome, tokens and trial in block. The plot shows the coefficients of each regressor and their %95 confidence interval in a mixed effects logistic (for switch) or mixed linear (for confidence) regression model, controlling for the full structure random effects of participant identity. Outcome is coded as a numerical variable with 'nNR' as n. Tokens is coded as numerical indicating the number of tokens captured. Trial in block is a numerical variable defined as the distance of the trial from the subjective start of the block. Condition is a 2-level factor, contrast coded with simple coding.
(B) Same as (A) for monkeys except the model is a simple logistic regression, combining data from two monkeys.

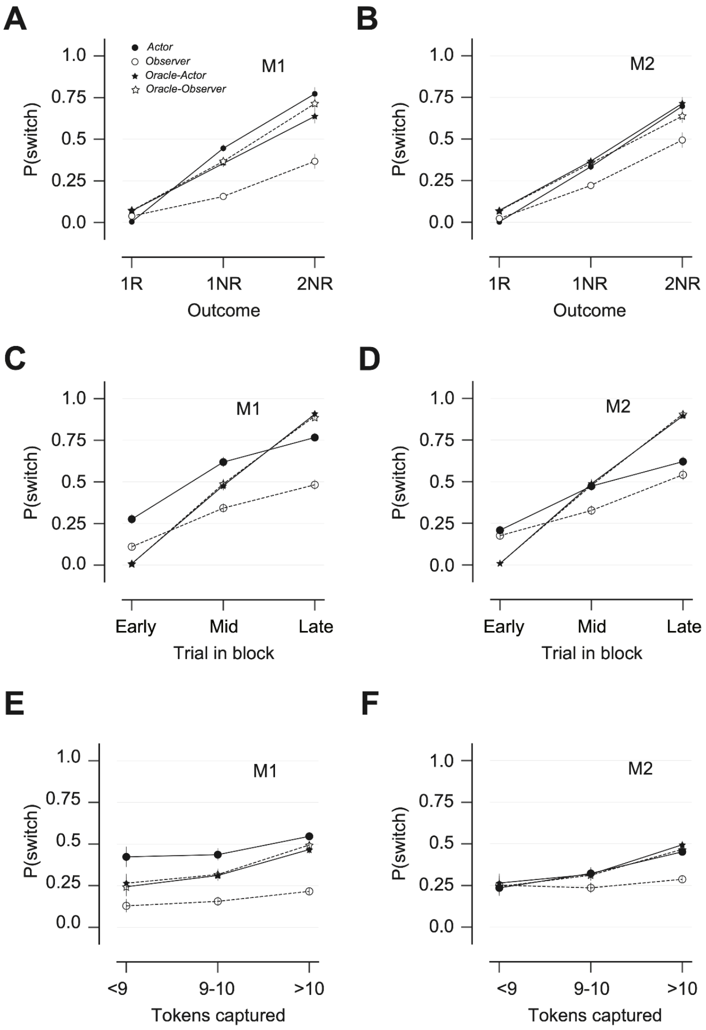


Figure S4. Effect of outcome, trial in block, and tokens captured on individual monkeys.

(A) Probability of switching to the other side (P(switch)) conditioned on trial outcome for monkey M1. Solid line with filled circle: actor trials. Dashed line with open circles: observer trials. Solid line with filled star: artificial oracle agent (methods) in actor trials. Dashed line with open star: artificial oracle agent in observer trials.

(B) Same as A for M2.

(C-D) Same as A-B for trial in block.

(E-F) Same as A-B for the number of tokens captured.

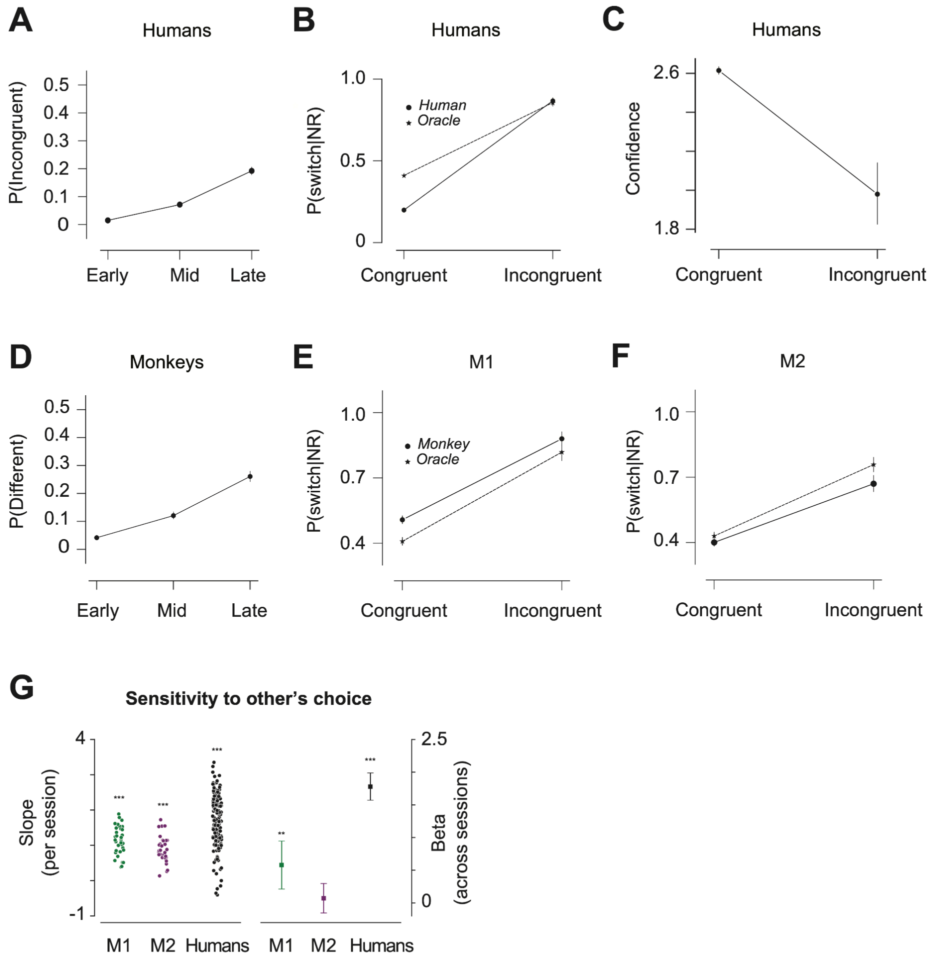


Figure S5. Effect of choice congruence on switching behavior and confidence

(A) Proportion of trials where the players make different choices as a function of trial in block. Early: trials 1-5; Mid: trials 6-10; Late: trials 11+. Data from all sessions in human participants.
(B) Proportion of trials where participants switched after an unrewarded trial as an actor, conditioned on trial congruence. Oracle: artificial oracle agent which switches on objective switch trials (see methods).
(C) Reported confidence pooled from human participants across all sessions, conditioned on trial congruence.
(D) Same as (A) for monkeys.
(E-F) Same as (B) for M1 and M2.
(G) Left: slopes of simple logistic regression for P(switch) over other's choice (congruent coded as 0, incongruent coded as 1), computed from actor trials for each participant for each session, where stars indicate P values from two-sided t test (***:P<0.001, **:P<0.01, *:P<0.05); Right: beta values from the full logistic regression for P(switch) over outcome, trial in block, number of captured tokens, and congruence, computed from actor trials for each participant over all

sessions, where vertical lines indicate 95% confidence interval, and stars indicate P values from two-sided t test (***:P<0.001, **:P<0.01, *:P<0.05).

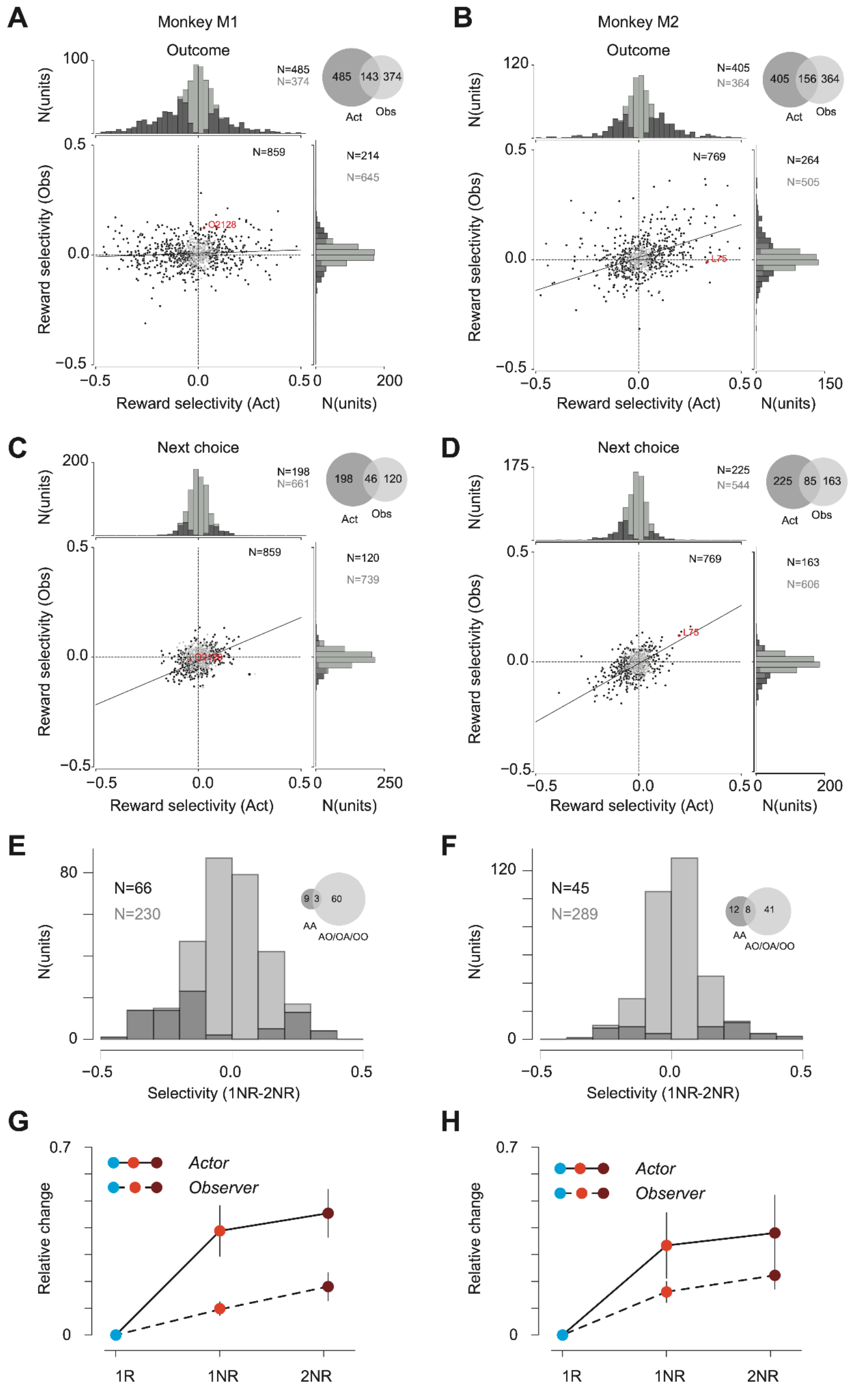


Figure S6. ACC neurons encode and integrate actor and observer outcome

(A) Reward selectivity in actor and observer conditions following outcome, including all units recorded from M1. Lower left: scatter plot of actor (x axis) and observer (y axis) reward selectivity, computed as the area under the curve of receiver operating characteristic (ROC) analysis based on spike count in the first 600 ms following outcome feedback. Straight line is the total regression performed on the selectivity values (regression slope: 0.03+/-0.02;

explained variance: 0.5%). Black dots are those neurons with significant selectivity in either condition. Green and magenta squares are example neurons from either animal. Top left: histogram of reward selectivity in the actor condition. Black bars correspond to neurons with significant selectivity (485/859; permutation test, 1000 times, P<0.05). Right: same histogram for the observer condition (significant neurons:214/859). Top right: number of neurons that are reward selective in the actor condition (485), observer condition (214), and both (143).

(B) Same as (A) for M2. Regression slope: 0.30+/-0.03; explained variance: 17.0%.

(C) Same as (A) but for the 600 ms before M1 makes choice contact. Regression slope: 0.40+/-0.04; explained variance: 26.0%.

(D) Same as (C) for M2. Regression slope: 0.53+/-0.04; explained variance: 36.0%.

(E) Histogram of accumulation selectivity defined as the area under the curve of ROC analysis based on spike count in the first 600 ms following outcome feedback, performed on the rate difference between 1NR vs. 2NR, separated into 1NR-Actor, 2NR-Actor (AA), AO, OA, OO conditions. Data from M1. Actor or observer reward selective neurons that share sign in reward selectivity are included in this analysis. Black bars correspond to neurons with significant selectivity in any of the four conditions (66/296; permutation test, 1000 times, P<0.05).

(F) Same as (E) for M2.

(G) Relative change in z-scored firing rate in the 600 ms window following outcome feedback between 1R (actor), 1NR (actor or observer), 2NR (actor or observer). Neurons from M1 with significant integration selectivity are included in this analysis. Solid line corresponds to actor conditions in 1NR and 2NR, dashed line corresponds to observer conditions. For neurons with decreased firing rate in 1NR vs. 1R, the rate change is multiplied by -1 for both 1NR and 2NR. Error bars indicate 95% confidence interval from bootstrap, N=1000; Actor-1NR: [0.30, 0.48]; Actor-2NR: [0.37, 0.54]; Observer-1NR: [0.08, 0.12]; Observer-2NR: [0.13, 0.23]. p<0.001 for comparing Actor to Observer conditions in both 1NR and 2NR.

(H) Same as (G) for M2. Confidence interval for Actor-1NR: [0.21, 0.45]; Actor-2NR: [0.25, 0.52]; Observer-1NR: [0.12, 0.21]; Observer-2NR: [0.17, 0.28]. p<0.01 for 1NR, p<0.05 for 2NR.

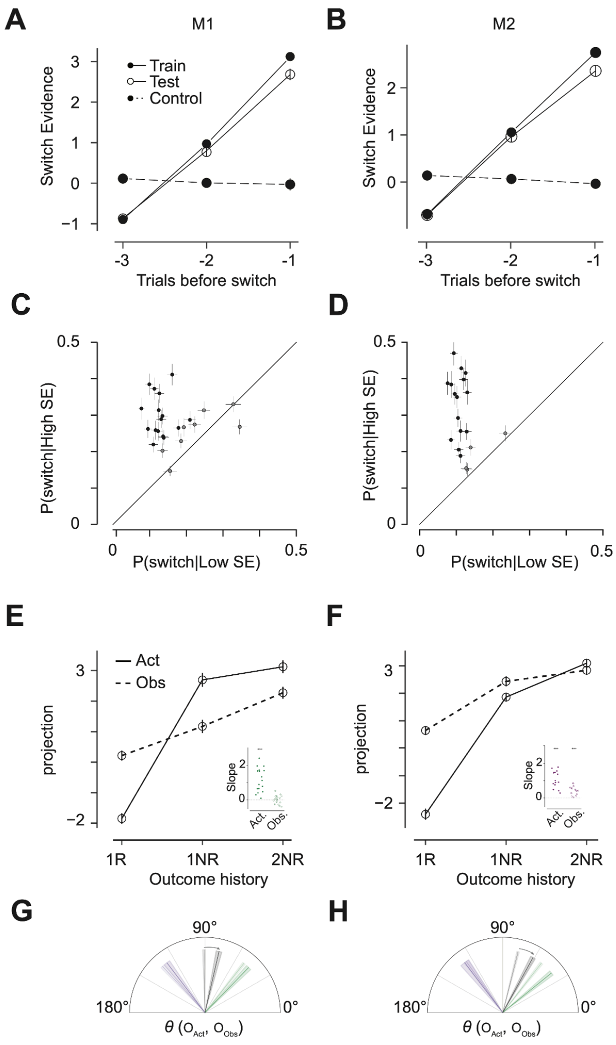


Figure S7. Population geometry of multi-agent evidence integration

(A) Projection on the identified switch dimension relative to number of trials before switch for the data used to compute the dimension (solid disks with solid line), held out test trials (open circles with solid line), and a null control which is selected by treating every three trials as 3,2,1 trials before switch, irrespective of actual switch trials. Data from M1 across sessions.
(B) Same as (A) for M2.
(C) Proportion of switch trials conditioned on a low (x axis) or high (y axis) projection on the switch evidence dimension for each session. Error bars: SEM. Black dots represent sessions with significant differences between low and high projection groups (N=16/24 sessions, rank sum test, P<0.05). Trials are sorted by the projection on switch evidence dimension and labeled as high or low SE based on the projection value greater or less than the median for each session.
(D) Same as (C) for M2. Significant sessions: 15/19.

(E) Projection on switch evidence dimension by behavioral condition. Data from M1. Solid line: actor conditions. Dashed line: observer conditions. Inset: slope of regression line for projection on SE over outcome history in actor and observer conditions.
(F) Same as (E) for M2.
(G) Angle between actor and observer outcome dimensions (black), and between aligned (green) or anti-aligned (magenta) subspaces. Lighter shades: angle computed from [0,600ms] window after outcome. Darker shade: computed from [-600,0ms] before choice. Data from M1.
(H) Same as (G) for M2.

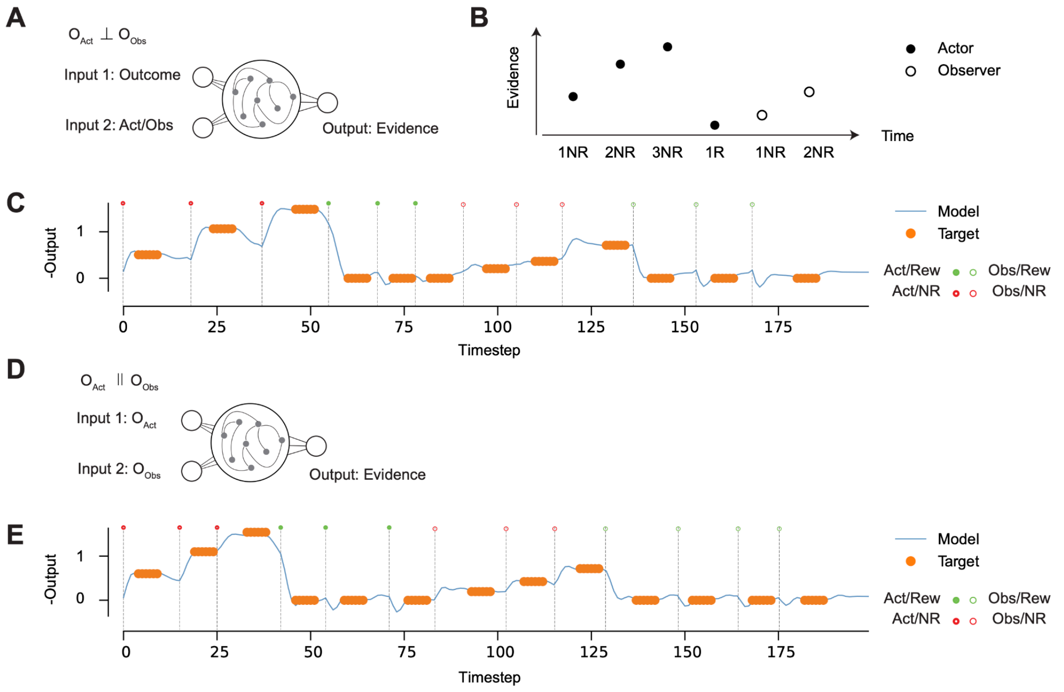


Figure S8. RNN architecture, task, and output

(A) Recurrent neural network instantiating the parallel hypothesis. Inputs represent the agent-agnostic outcome and indicator variable for actor or observer conditions. A trainable RNN in the center learns to generate an output that is the integrated evidence.
(B) Illustration of trial sequence and training target output. X axis represent six trials during training which can be categorized as 1R trials (where outcome is rewarded) and nNR trials (where outcome is unrewarded, consecutive for n trials). Y axis represent the desired outcome for each trial, which increments for every unrewarded trial, with higher increase for actor (filled circles) vs. observer (open circles) trials, and resets to zero for rewarded trials. Note in this simplified task there is no choice phase, and the network only reports integrated evidence.
(C) Example model output in a test session. X axis represents time steps. Y axis represents the inverse of model output (blue line) and target output (orange circles), which the model is evaluated against. Note the model output is only evaluated at target time. The output is shown in inverse because the model output is the weighted sum of recent negative prediction errors, which is the inverse of cumulative evidence for a decision to switch (not implemented). Vertical dashed lines represent the input time for each trial. Circles on top of the vertical lines represent trial type. Filled circles: actor trials. Open circles: observer trials. Red: unrewarded trials. Green: rewarded trials.
(D) Same as (A) for the orthogonal hypothesis. Inputs each represent the actor and observer outcome.
(E) Same as (C) for the network in (D).

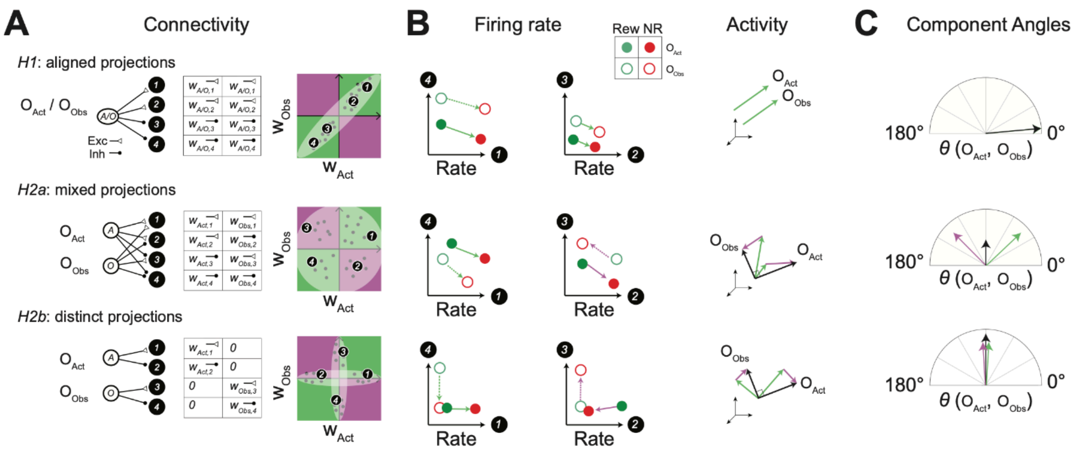


Figure S9. Angle decomposition analysis.

(A) Left: hypothesized connectivity patterns between inputs representing actor and/or observer outcome and integrator network. Each row illustrates the idealized input node as circles on the left, with net effective excitatory or inhibitory connections to receiving units in the network (black disks). Middle: weight matrix for each connectivity pattern, where each row is a unit, left column is the effective weight from actor outcomes (unrewarded vs. rewarded trials), and right column is the effective weight from observer outcomes. Right: visualization of the weight matrix in two dimensions, where x axis is the weight from actor outcome and y axis is the weight from observer outcome. Each dot represents a unit, with units 1-4 labeled. This space is separated into the positively correlated subspace (‘aligned subspace’, green) and negatively correlated (‘anti-aligned subspace’, magenta).

(B) Left: firing rate of the labeled units from (A). Each axis represents the mean firing rate of the unit identified at the end (1-4), and each circle represents the mean firing rate of that pair under one condition. Green circles: rewarded trials. Red circles: unrewarded trials. Filled circles: actor trials. Open circles: observer trials. Arrows point from reward trials to unrewarded trials, which represents the subcomponent of actor (solid arrows) or observer (dashed arrows) outcome dimensions. Color of arrows indicate which subspace (green: aligned; magenta: anti-aligned) both units are sampled from. Right: outcome dimensions in actor ($O_{act}$) and observer ($O_{Obs}$) conditions. Green arrows represent the subcomponent of each outcome dimension constituted by units in the aligned subspace, and magenta arrows represent those from units in the anti-aligned subspace.

(C) Component angles between $O_{act}$ and $O_{Obs}$. Black arrows: angle between full outcome dimensions. Green: angle between aligned subcomponents. Magenta: angle between anti-aligned subcomponents. Note in the top row, green and black vectors overlap because all units are in the aligned subspace, and there is no magenta arrow because there are no units in the anti-aligned subspace.

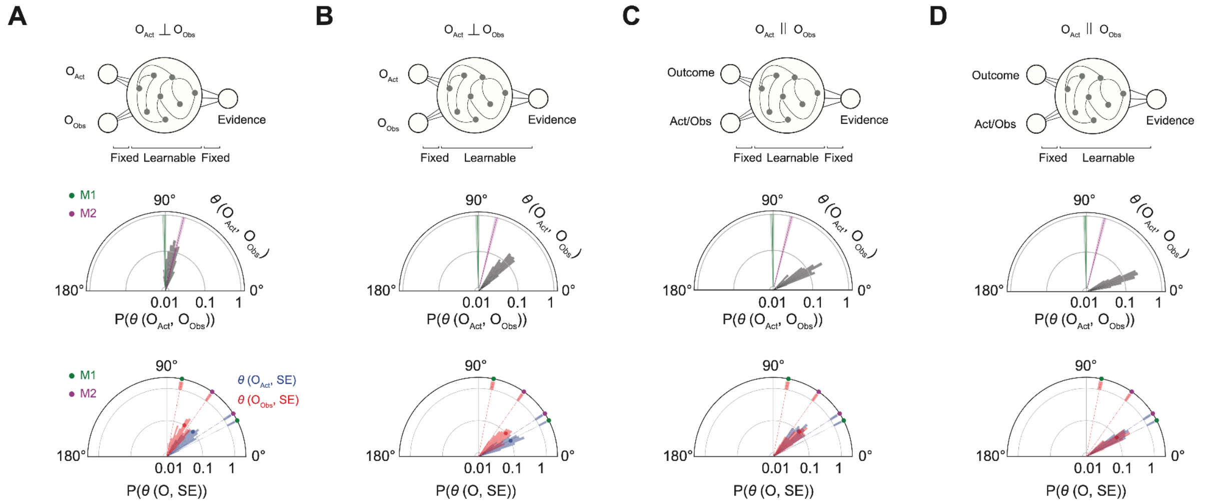


Figure S10. Readout-fixed and readout-learnable networks

(A) Top: Readout-fixed orthogonal networks, same as in Figure 4J. Fixed refers to those weights that were not subject to training updates. Learnable refers to the other parameters that were updated during training. Middle: comparison of actor/observer outcome angles in networks and ACC data in monkeys, same as Figure 4L. Bottom: Comparison of outcome/switch evidence angles in networks and ACC data in monkeys, same as Figure 4P. $\theta(O_{Act}, O_{Obs})$:78.88 ± 7.56; $\theta(O_{Act}, SE)$: 43.15 ± 10.22; $\theta(O_{Obs}, SE)$: 59.11±10.89 degrees.
(B) Same as A for readout-learnable orthogonal networks. $\theta(O_{Act}, O_{Obs})$: 45.91 ± 6.26; $\theta(O_{Act}, SE)$: 26.04 ± 4.33, $\theta(O_{Obs}, SE)$: 40.18 ± 7.61 degrees.
(C) Same as A for readout-fixed parallel networks. $\theta(O_{Act}, O_{Obs})$: 30.77 ± 6.88; $\theta(O_{Act}, SE)$: 44.03 ± 7.35, $\theta(O_{Obs}, SE)$: 43.48 ± 7.80 degrees.
(D) Same as A for readout-learnable parallel networks. $\theta(O_{Act}, O_{Obs})$: 20.92 ± 5.48; $\theta(O_{Act}, SE)$: 31.58 ± 5.39, $\theta(O_{Obs}, SE)$: 31.79 ± 5.78 degrees.

Table S1. ACC recording location. ML: mediolateral coordinate relative to the midline. AG: anterior-posterior coordinate relative to the genu of the arcuate sulcus.

| | Monkey 1 | Monkey 2 |
|---|---|---|
| Stereotactic coordinates | ML 2-5mm | ML 1-5 mm |
| | AG 2-11mm | AG 2-11 mm |
| Recording sessions/trials | 31/29917 | 20/19196 |
